## Supplementary material for "Data-Driven Modeling and Analysis of Fatty Acid Desaturase in Plants": TeX Files: paper.bib

@article{pfam,
    author = {Mistry, Jaina and Chuguransky, Sara and Williams, Lowri and Qureshi, Matloob and Salazar, Gustavo A and Sonnhammer, Erik L L and Tosatto, Silvio C E and Paladin, Lisanna and Raj, Shriya and Richardson, Lorna J and Finn, Robert D and Bateman, Alex},
    title = "{Pfam: The protein families database in 2021}",
    journal = {Nucleic Acids Research},
    volume = {49},
    number = {D1},
    pages = {D412-D419},
    year = {2020},
    month = {10},
    abstract = "{The Pfam database is a widely used resource for classifying protein sequences into families and domains. Since Pfam was last described in this journal, over 350 new families have been added in Pfam 33.1 and numerous improvements have been made to existing entries. To facilitate research on COVID-19, we have revised the Pfam entries that cover the SARS-CoV-2 proteome, and built new entries for regions that were not covered by Pfam. We have reintroduced Pfam-B which provides an automatically generated supplement to Pfam and contains 136 730 novel clusters of sequences that are not yet matched by a Pfam family. The new Pfam-B is based on a clustering by the MMseqs2 software. We have compared all of the regions in the RepeatsDB to those in Pfam and have started to use the results to build and refine Pfam repeat families. Pfam is freely available for browsing and download at http://pfam.xfam.org/.}",
    issn = {0305-1048},
    doi = {10.1093/nar/gkaa913}
}
@article{Fox2004,
abstract = {Stearoyl-acyl carrier protein ∆ 9 desaturase (∆9D) produces oleic acid, a nutritionally valuable fatty acid containing a cis double bond between C-9 and C-10. This multiprotein diiron enzyme complex reacts with stearoyl-acyl carrier protein, reduced [2Fe-2S] ferre-doxin, and O 2 to complete the highly regiospecific and stereo-selective desaturation reaction. Interactions with the acyl chain provide stability to the enzyme-substrate complex, give an energetic contribution to catalytic selectivity, and help to order the electron transfer, O 2 binding, and C-H bond cleavage steps of catalysis. Reactions with natural acyl chains indicate the involvement of a highly reactive diiron intermediate capable of oxidizing secondary C-H bonds (bond dissociation energy ≈ 95 kcal/mol), but also capable of diagnostic O-atom transfer reactions with the appropriate substrate analogues. For soluble ∆9D, the natural reaction may initiate at the C-10 position, in contrast to the well-established initial reactivity of the membrane enzyme homologue stearoyl-coenzyme A (CoA) ∆ 9 desaturase at the C-9 position.},
author = {Fox, Brian G and Lyle, Karen S and Rogge, Corina E},
doi = {10.1021/ar030186h},
title = {{Reactions of the Diiron Enzyme Stearoyl-Acyl Carrier Protein Desaturase}},
year = {2004}
}
@article {Schmidt9477,
	author = {Schmidt, H and Heinz, E},
	title = {Desaturation of oleoyl groups in envelope membranes from spinach chloroplasts.},
	volume = {87},
	number = {23},
	pages = {9477--9480},
	year = {1990},
	doi = {10.1073/pnas.87.23.9477},
	publisher = {National Academy of Sciences},
	abstract = {Envelope membranes isolated from chloroplasts of spinach (Spinacia oleracea) desaturate oleoyl groups in monogalactosyl diacylglycerol to linoleoyl groups. The desaturation requires NADPH in combination with ferredoxin and is not restricted to monogalactosyl diacylglycerol, since it is also observed in biosynthetic intermediates as, for example, in phosphatidic acid. This indicates a certain degree of unspecificity of the oleate desaturase in isolated envelope membranes. Lipid desaturation is another important function of chloroplast envelopes.},
	issn = {0027-8424},
	journal = {Proceedings of the National Academy of Sciences}
}
@article{Schmidt1990,
    author = {Schmidt, Hermann and Heinz, Ernst},
    title = "{Involvement of Ferredoxin in Desaturation of Lipid-Bound Oleate in Chloroplasts 1}",
    journal = {Plant Physiology},
    volume = {94},
    number = {1},
    pages = {214-220},
    year = {1990},
    month = {09},
    abstract = "{Intact spinach (Spinacia oleracea) chloroplasts, pulse-labeled with [14C]acetate, desaturate newly formed fatty acids as ester groups of monogalactosyl diacylglycerol in a subsequent chase in the dark. Rupture of pulse-labeled chloroplasts by addition of a detergent solution 3-([3-cholamidopropyl]dimethylammonio)-1-propane sulfonate preserves part of this desaturation activity. Direct addition of different free fatty acids together with appropriate cofactors to detergent-ruptured chloroplasts results in fatty acid labeling of monogalactosyl diacylglycerol. During subsequent incubation these lipid-linked fatty acids are desaturated, i.e. 18:1 to 18:2 and 18:3 and to a small extent also 16:0 to 16:3. The formation of 18:2 was also observed after incorporation of 18:1 into sulfolipid and phosphatidyl choline. Density gradient centrifugation separated a membrane fraction from detergent-ruptured chloroplasts which in the presence of appropriate cofactors incorporated 18:1 and 18:2 into the above-mentioned lipids. In the light, desaturation was dependent on added ferredoxin, whereas in the dark, in addition to ferredoxin NAD(P)H was also required. Preliminary evidence for the involvement of membrane-bound ferredoxin:NADP oxidoreductase (FNR) as a third component of desaturation in the dark was obtained by inhibitor studies including antibodies against FNR. Desaturation of lipid-bound 18:1 and 18:2 resembles stearoyl-ACP desaturation with respect to its requirement of reduced ferredoxin and oxygen.}",
    issn = {0032-0889},
    doi = {10.1104/pp.94.1.214},
}
@article{Smith1990,
    author = {Smith, M A and Cross, A R and Jones, O T G and Griffiths, W T and Stymne, S and Stobart, K},
    title = "{Electron-transport components of the 1-acyl-2-oleoyl-sn-glycero-3-phosphocholine {$\Delta$}12-desaturase ({$\Delta$}12-desaturase) in microsomal preparations from developing safflower (Carthamus tinctorius L.) cotyledons}",
    journal = {Biochemical Journal},
    volume = {272},
    number = {1},
    pages = {23-29},
    year = {1990},
    month = {11},
    abstract = "{The major cytochrome in microsomal membrane preparations from developing seeds of safflower (Carthamus tinctorius, var High Linoleate), has a reduced-minus-oxidized difference spectrum characteristic of a b-type cytochrome, and was identified from its midpoint-potential (E'7.2) value as cytochrome b5. Cytochromes P-450 and P-420 were also present. The cytochrome b5 content of microsomal preparations from a number of oilseed species was found to be in the order of 200-300 pmol/mg of protein. The cytochrome b5 was reduced in the membrane preparations by NADH, demonstrating the presence of an NADH: cytochrome b5 reductase; NADPH was a less effective donor. Microsomal membranes catalysed the NAD(P)H-dependent conversion of radioactive oleate into linoleate, indicating acyl-CoA: lysophosphatidylcholine acyltransferase and 1-acyl-2-oleoyl-sn-glycero-3-phosphocholine delta 12-desaturase (delta 12-desaturase) activity. Desaturation of oleate to linoleate was unaffected by CO, but inhibited by CN-. The addition of oleoyl-CoA to the NADH-reduced membranes resulted in the CN(-)-sensitive partial re-oxidation of cytochrome b5, indicating that electrons from NADH were transferred to the site of desaturation via this cytochrome. The delta 12-desaturase in safflower, therefore, is CN(-)-sensitive and appears to require cytochrome b5 and NADH: cytochrome b5 reductase for activity.}",
    issn = {0264-6021},
    doi = {10.1042/bj2720023},
}
@article{KEARNS1991431,
title = {The role of cytochrome b5 in {$\Delta$}12 desaturation of oleic acid by microsomes of safflower (Carthamus tinctorius L.)},
journal = {Archives of Biochemistry and Biophysics},
volume = {284},
number = {2},
pages = {431-436},
year = {1991},
issn = {0003-9861},
doi = {10.1016/0003-9861(91)90319-E},
author = {Ellen V. Kearns and Suzanne Hugly and Chris R. Somerville},
abstract = {The electron donors for the membrane-bound fatty acid desaturases of higher plants have not previously been identified. In order to assess the participation of cytochrome b5 in microsomal fatty acid desaturation, the cytoplasmic domain of microsomal cytochrome b5 was purified from Brassica oleracea, and murine polyclonal antibodies were prepared. The IgG fraction from ascites fluid inhibited 62\% of NADH-dependent cytochrome c reduction in safflower (Carthamus tinctorius L.) microsomes. These antibodies also blocked desaturation of oleic acid to linoleic acid in lipids of C. tinctorius microsomes by 93\%, suggesting that cytochrome b5 is the electron donor for the $\Delta$12 desaturase.}
}
@article{Falcone1994,
    author = {Falcone, D. L. and Gibson, S. and Lemieux, B. and Somerville, C.},
    title = "{Identification of a Gene that Complements an Arabidopsis Mutant Deficient in Chloroplast [omega]6 Desaturase Activity}",
    journal = {Plant Physiology},
    volume = {106},
    number = {4},
    pages = {1453-1459},
    year = {1994},
    month = {12},
    abstract = "{Membrane lipids of the fad6 (formerly fadC) mutant of Arabidopsis, which is deficient in chloroplast [omega]6 desaturase activity, have increased levels of monounsaturated fatty acids and are deficient in trienoic fatty acids. A putative fad6 cDNA clone was isolated by probing a cDNA library with a degenerate oligonucleotide based on a conserved region within known [omega]3 desaturase genes. Expression of the cDNA in transgenic plants of a fad6 mutant restored normal levels of all fatty acids. When used as a hybridization probe, the cDNA identified a restriction fragment-length polymorphism that co-segregated with the fad6 mutation. Thus, on the basis of a genetic complementation test and genetic map position, the fad6 gene is encoded by the cDNA. The cDNA encoded a 418-amino acid polypeptide of 47,727 D that displayed a high degree of sequence similarity to a [delta]12 desaturase from the cyanobacterium Synechocystis. The fad6 gene exhibited less sequence homology to any known higher plant desaturase, including an endoplasmic reticulum-localized [omega]6 desaturase corresponding to the Arabidopsis fad2 gene.}",
    issn = {0032-0889},
    doi = {10.1104/pp.106.4.1453},
}
@article{Sperling2001,
author = {Sperling, Petra and Heinz, Ernst},
title = {Desaturases fused to their electron donor},
journal = {European Journal of Lipid Science and Technology},
volume = {103},
number = {3},
pages = {158-180},
keywords = {cytochrome b5 domain, front-end desaturases, long-chain polyunsaturated fatty acids},
doi = {10.1002/1438-9312(200103)103:3<158::AID-EJLT158>3.0.CO;2-1},
abstract = {Abstract Fatty acid desaturations in the carboxy-terminal segment from C1—C10 are catalyzed in many, but not in all cases, by desaturase enzymes which are fused to their electron donor cytochrome b5. Several of these enzymes (“front-end desaturases”) from a wide variety of organisms have been cloned and functionally expressed for proof of regio-, stereo- and chain length-selectivity. In most cases the actual status of the substrate chain, whether coenzyme A thioester or component of a membrane lipid, is not known. The cytochrome b5 domain is located N-terminally, internally or C-terminally. Compared to the free cytochrome b5 , the fused domains show a significant reduction of acidic amino acid residues on the surface of the four helices enclosing the heme group. It is discussed how this may contribute to hydrophobic domain pairing required for interdomain electron transport. This is in contrast to the mode of interaction of free cytochrome b5 with its partners, which is governed by electrostatic charge pairing. A look at crystallized or computer-simulated models involving fused or free cytochrome b5 helps to outline the problems encountered by optimizing the docking of partners and the exchange of electrons between domains of different degrees of mobility.},
year = {2001}
}
@article{LI2016341,
title = {Classification and substrate head-group specificity of membrane fatty acid desaturases},
journal = {Computational and Structural Biotechnology Journal},
volume = {14},
pages = {341-349},
year = {2016},
issn = {2001-0370},
doi = {10.1016/j.csbj.2016.08.003},
author = {Dongdi Li and Ruth Moorman and Thomas Vanhercke and James Petrie and Surinder Singh and Colin J. Jackson},
keywords = {Membrane-bound fatty acid desaturases, Lipid-substrate specificity, Sequence similarity networks (SSNs)},
abstract = {Membrane fatty acid desaturases are a diverse superfamily of enzymes that catalyze the introduction of double bonds into fatty acids. They are essential in a range of metabolic processes, such as the production of omega-3 fatty acids. However, our structure–function understanding of this superfamily is still developing and their range of activities and substrate specificities are broad, and often overlapping, which has made their systematic characterization challenging. A central issue with characterizing these proteins has been the lack of a structural model, which has been overcome with the recent publication of the crystal structures of two mammalian fatty acid desaturases. In this work, we have used sequence similarity networks to investigate the similarity among over 5000 related membrane fatty acid desaturase sequences, leading to a detailed classification of the superfamily, families and subfamilies with regard to their function and substrate head-group specificity. This work will facilitate rapid prediction of the function and specificity of new and existing sequences, as well as forming a basis for future efforts to manipulate the substrate specificity of these proteins for biotechnology applications.}
}
@article{Bai2015,
author = {Bai, Yonghong and McCoy, Jason G. and Levin, Elena J. and Sobrado, Pablo and Rajashankar, Kanagalaghatta R. and Fox, Brian G. and Zhou, Ming},
doi = {10.1038/nature14549},
issn = {14764687},
journal = {Nature},
month = {aug},
number = {7564},
pages = {252--256},
pmid = {26098370},
publisher = {Nature Publishing Group},
title = {{X-ray structure of a mammalian stearoyl-CoA desaturase}},
volume = {524},
year = {2015}
}
@article{Tocher1998,
author = {Tocher, D.R. and Leaver, M.J. and Hodgson, P.A.},
doi = {10.1016/S0163-7827(98)00005-8},
issn = {01637827},
journal = {Progress in Lipid Research},
month = {jul},
number = {2-3},
pages = {73--117},
title = {{Recent advances in the biochemistry and molecular biology of fatty acyl desaturases}},
volume = {37},
year = {1998}
}
@article{Reed2000,
author = {Reed, Darwin W. and Sch{\"{a}}fer, Ulrike A. and Covello, Patrick S.},
doi = {10.1104/pp.122.3.715},
issn = {0032-0889},
journal = {Plant Physiology},
month = {mar},
number = {3},
pages = {715--720},
title = {{Characterization of the Brassica napus Extraplastidial Linoleate Desaturase by Expression in Saccharomyces cerevisiae}},
volume = {122},
year = {2000}
}
@misc{Meesapyodsuk2012,
abstract = {Very long chain polyunsaturated fatty acids such as arachidonic acid (ARA, 20:4n-6), eicosapentaenoic acid (EPA, 20:5n-3), docosapentaenoic acid (DPA, 22:5n-3) and docosahexaenoic acid (DHA, 22:6n-3) are essential components of cell membranes, and are precursors for a group of hormone-like bioactive compounds (eicosanoids and docosanoids) involved in regulation of various physiological activities in animals and humans. The biosynthesis of these fatty acids involves an alternating process of fatty acid desaturation and elongation. The desaturation is catalyzed by a unique class of oxygenases called front-end desaturases that introduce double bonds between the preexisting double bond and the carboxyl end of polyunsaturated fatty acids. The first gene encoding a front-end desaturase was cloned in 1993 from cyanobacteria. Since then, front-end desaturases have been identified and characterized from a wide range of eukaryotic species including algae, protozoa, fungi, plants and animals including humans. Unlike front-end desaturases from bacteria, those from eukaryotes are structurally characterized by the presence of an N-terminal cytochrome b5-like domain fused to the main desaturation domain. Understanding the structure, function and evolution of front-end desaturases, as well as their roles in the biosynthesis of very long chain polyunsaturated fatty acids offers the opportunity to engineer production of these fatty acids in transgenic oilseed plants for nutraceutical markets. {\textcopyright} AOCS 2011.},
author = {Meesapyodsuk, Dauenpen and Qiu, Xiao},
booktitle = {Lipids},
doi = {10.1007/s11745-011-3617-2},
issn = {00244201},
keywords = {Biotechnology,Desaturases,General area,Lipid biochemistry,N-3 fatty acids,N-6 fatty acids,Polyunsaturated fatty acids (PUFA)},
month = {mar},
number = {3},
pages = {227--237},
pmid = {22009657},
publisher = {John Wiley {\&} Sons, Ltd},
title = {{The front-end desaturase: Structure, function, evolution and biotechnological use}},
volume = {47},
year = {2012}
}
@article{Yadav1993,
author = {Yadav, N. S. and Wierzbicki, A. and Aegerter, M. and Caster, C. S. and Perez-Grau, L. and Kinney, A. J. and Hitz, W. D. and {Booth Jr.}, J. R. and Schweiger, B. and Stecca, K. L. and Allen, S. M. and Blackwell, M. and Reiter, R. S. and Carlson, T. J. and Russell, S. H. and Feldmann, K. A. and Pierce, J. and Browse, J.},
doi = {10.1104/pp.103.2.467},
issn = {0032-0889},
journal = {Plant Physiology},
month = {oct},
number = {2},
pages = {467--476},
title = {{Cloning of Higher Plant $\omega$-3 Fatty Acid Desaturases}},
volume = {103},
year = {1993}
}
@article{Spychalla1997,
author = {Spychalla, J. P. and Kinney, A. J. and Browse, J.},
doi = {10.1073/pnas.94.4.1142},
issn = {0027-8424},
journal = {Proceedings of the National Academy of Sciences},
month = {feb},
number = {4},
pages = {1142--1147},
title = {{Identification of an animal -3 fatty acid desaturase by heterologous expression in Arabidopsis}},
volume = {94},
year = {1997}
}
@article{Meesapyodsuk2000,
author = {Meesapyodsuk, Dauenpen and Reed, Darwin W. and Savile, Christopher K. and Buist, Peter H. and Ambrose, Stephen J. and Covello, Patrick S.},
doi = {10.1021/bi000756a},
issn = {0006-2960},
journal = {Biochemistry},
month = {oct},
number = {39},
pages = {11948--11954},
title = {{Characterization of the Regiochemistry and Cryptoregiochemistry of a Caenorhabditis elegans Fatty Acid Desaturase ( FAT-1 ) Expressed in Saccharomyces cerevisiae †}},
volume = {39},
year = {2000}
}
@article{Buist2004,
author = {Buist, Peter H.},
doi = {10.1039/b302094k},
issn = {0265-0568},
journal = {Natural Product Reports},
number = {2},
pages = {249},
title = {{Fatty acid desaturases: selecting the dehydrogenation channel}},
volume = {21},
year = {2004}
}
@article{Hitz1994,
author = {Hitz, W. D. and Carlson, T. J. and {Booth Jr}, J. R. and Kinney, A. J. and Stecca, K. L. and Yadav, N. S.},
doi = {10.1104/pp.105.2.635},
issn = {0032-0889},
journal = {Plant Physiology},
month = {jun},
number = {2},
pages = {635--641},
title = {{Cloning of a Higher-Plant Plastid [omega]-6 Fatty Acid Desaturase cDNA and Its Expression in a Cyanobacterium}},
volume = {105},
year = {1994}
}
@article{Tsai2014,
author = {Tsai, Chung-Jung and Nussinov, Ruth},
doi = {10.1371/journal.pcbi.1003394},
editor = {Livesay, Dennis R.},
issn = {1553-7358},
journal = {PLoS Computational Biology},
month = {feb},
number = {2},
pages = {e1003394},
title = {{A Unified View of “How Allostery Works”}},
volume = {10},
year = {2014}
}
@article{Kondakova2015,
author = {Kondakova, Tatiana and D'Heyg{\`{e}}re, Fran{\c{c}}ois and Feuilloley, Marc J. and Orange, Nicole and Heipieper, Hermann J. and {Duclairoir Poc}, C{\'{e}}cile},
doi = {10.1016/j.chemphyslip.2015.06.006},
issn = {00093084},
journal = {Chemistry and Physics of Lipids},
month = {sep},
pages = {27--42},
title = {{Glycerophospholipid synthesis and functions in Pseudomonas}},
volume = {190},
year = {2015}
}
@article{Liu2011,
abstract = {Acyl-CoA:diacylglycerol acyltransferase (EC 2.3.1.20) is a membrane protein present mainly in the endoplasmic reticulum. It catalyzes the final and committed step in the biosynthesis of triacylglycerol, which is the principal repository of fatty acids for energy utilization and membrane formation. Two distinct family members of acyl-CoA:diacylglycerol acyltransferase, known as DGAT1 and DGAT2, have been characterized in different organisms, including mammals, fungi, and plants. In this study, we characterized the functional role and topological orientation of signature motifs in yeast (Saccharomyces cerevisiae) DGAT2 using mutagenesis in conjunction with chemical modification. Our data provide evidence that both the N and C termini are oriented toward the cytosol and have different catalytic roles. A highly conserved motif, 129YFP131, and a hydrophilic segment exclusive to yeast DGAT2 reside in a long endoplasmic reticulum luminal loop following the first transmembrane domain and play an essential role in enzyme catalysis. In addition, the strongly conserved His195 within the motif HPHG, which may play a role in the active site of DGAT2, is likely embedded in the membrane. These results indicate some similarities to the topology model of murine DGAT2 but also reveal striking differences suggesting that the topological organization of DGAT2 is not ubiquitously conserved. {\textcopyright} 2011 by The American Society for Biochemistry and Molecular Biology, Inc.},
author = {Liu, Qin and Siloto, Rodrigo M.P. and Snyder, Crystal L. and Weselake, Randall J.},
doi = {10.1074/jbc.M110.204412},
issn = {00219258},
journal = {Journal of Biological Chemistry},
month = {apr},
number = {15},
pages = {13115--13126},
pmid = {21321129},
publisher = {Elsevier},
title = {{Functional and topological analysis of yeast acyl-CoA:diacylglycerol acyltransferase 2, an endoplasmic reticulum enzyme essential for triacylglycerol biosynthesis}},
volume = {286},
year = {2011}
}
@article{Aranko2013,
author = {Aranko, A Sesilja and Oeemig, Jesper S and Kajander, Tommi and Iwa{\"{i}}, Hideo},
doi = {10.1038/nchembio.1320},
issn = {1552-4450},
journal = {Nature Chemical Biology},
month = {oct},
number = {10},
pages = {616--622},
title = {{Intermolecular domain swapping induces intein-mediated protein alternative splicing}},
volume = {9},
year = {2013}
}
@article{Shanklin1998,
author = {Shanklin, John and Cahoon, Edgar B.},
doi = {10.1146/annurev.arplant.49.1.611},
issn = {1040-2519},
journal = {Annual Review of Plant Physiology and Plant Molecular Biology},
month = {jun},
number = {1},
pages = {611--641},
title = {{DESATURATION AND RELATED MODIFICATIONS OF FATTY ACIDS}},
volume = {49},
year = {1998}
}
@article{Cahoon2006,
abstract = {Expression of $\Delta$12-oleic acid desaturase-related fatty acid conjugases from Calendula officinalis, Momordica charantia, and Vernicia fordii in seeds of soybean (Glycine max) or an Arabidopsis thaliana fad3/fae1 mutant was accompanied by the accumulation of the conjugated fatty acids calendic acid or $\alpha$-eleostearic acid to amounts as high as 20{\%} of the total fatty acids. Conjugated fatty acids, which are synthesized from phosphatidylcholine (PC)-linked substrates, accumulated in PC and phosphatidylethanolamine, and relative amounts of these fatty acids were higher in PC than in triacylglycerol (TAG) in the transgenic seeds. The highest relative amounts of conjugated fatty acids were detected in PC from seeds of soybean and A. thaliana that expressed the C. officinalis and M. charantia conjugases, where they accounted for nearly 25{\%} of the fatty acids of this lipid class. In these seeds, {\textgreater}85{\%} of the conjugated fatty acids in PC were detected in the sn-2 position, and these fatty acids were also enriched in the sn-2 position of TAG. In marked contrast to the transgenic seeds, conjugated fatty acids composed {\textless}1.5{\%} of the fatty acids in PC from seeds of five unrelated species that naturally synthesize a variety of conjugated fatty acid isomers, including seeds that accumulate conjugated fatty acids to {\textgreater}80{\%} of the total fatty acids. These results suggest that soybean and A. thaliana seeds are deficient in their metabolic capacity to selectively catalyze the flux of conjugated fatty acids from their site of synthesis on PC to storage in TAG.},
author = {Cahoon, Edgar B and Dietrich, Charles R and Meyer, Knut and Damude, Howard G and Dyer, John M and Kinney, Anthony J},
doi = {10.1016/j.phytochem.2006.04.013},
issn = {0031-9422},
journal = {Phytochemistry},
keywords = {Calendic acid,Conjugated fatty acid,FAD2,Fatty acid conjugase,Leguminosae,Metabolic engineering,Oilseed engineering,Phosphatidylcholine,Soybean,Triacylglycerol,Unusual fatty acid,$\alpha$-Eleostearic acid},
number = {12},
pages = {1166--1176},
title = {{Conjugated fatty acids accumulate to high levels in phospholipids of metabolically engineered soybean and Arabidopsis seeds}},
volume = {67},
year = {2006}
}
@article{Caspi2012,
author = {Caspi, R. and Altman, T. and Dreher, K. and Fulcher, C. A. and Subhraveti, P. and Keseler, I. M. and Kothari, A. and Krummenacker, M. and Latendresse, M. and Mueller, L. A. and Ong, Q. and Paley, S. and Pujar, A. and Shearer, A. G. and Travers, M. and Weerasinghe, D. and Zhang, P. and Karp, P. D.},
doi = {10.1093/nar/gkr1014},
issn = {0305-1048},
journal = {Nucleic Acids Research},
month = {jan},
number = {D1},
pages = {D742--D753},
title = {{The MetaCyc database of metabolic pathways and enzymes and the BioCyc collection of pathway/genome databases}},
volume = {40},
year = {2012}
}
@article{Sperling2003,
author = {Sperling, P. and Ternes, P. and Zank, T. K. and Heinz, E.},
doi = {10.1016/S0952-3278(02)00258-2},
issn = {09523278},
journal = {Prostaglandins Leukotrienes and Essential Fatty Acids},
month = {feb},
number = {2},
pages = {73--95},
pmid = {12538072},
publisher = {Churchill Livingstone},
title = {{The evolution of desaturases}},
volume = {68},
year = {2003}
}
@article{Broadwater1998,
author = {Broadwater, John A. and Haas, Jeffrey A. and Fox, Brian G.},
doi = {10.1002/(SICI)1521-4133(19985)100:4/5<103::AID-LIPI103>3.0.CO;2-4},
issn = {0931-5985},
journal = {Lipid - Fett},
month = {may},
number = {4-5},
pages = {103--113},
title = {{The fundamental, versatile role of diiron enzymes in lipid metabolism}},
volume = {100},
year = {1998}
}
@article{McFie2010,
abstract = {Acyl CoA:diacylglycerol acyltransferase (DGAT) is an integral membrane protein of the endoplasmic reticulum that catalyzes the synthesis of triacylglycerols. Two DGAT enzymes have been identified (DGAT1 and DGAT2) with unique roles in lipid metabolism. DGAT1 is a multifunctional acyltransferase capable of synthesizing diacylglycerol, retinyl, and wax esters in addition to triacylglycerol. Here, we report the membrane topology for murine DGAT1 using protease protections assays and indirect immunofluorescence in conjunction with selective permeabilization of cellular membranes. Topology models based on prediction algorithms suggested that DGAT1 had eight transmembrane domains. In contrast, our data indicate that DGAT1 has three transmembrane domains with the N terminus oriented toward the cytosol. The C-terminal region of DGAT1, which accounts for ∼50{\%} of the protein, is present in the endoplasmic reticulum lumen and contains a highly conserved histidine residue (His-426) that may be part of the active site. Mutagenesis of His-426 to alanine impaired the ability of DGAT1 to synthesize triacylglycerols as well as retinyl and wax esters in an in vitro acyltransferase assay. Finally, we show that the N-terminal domain of DGAT1 is not required for the catalytic activity of DGAT1 but, instead, may be involved in regulating enzyme activity and dimer/tetramer formation. {\textcopyright} 2010 by The American Society for Biochemistry and Molecular Biology, Inc.},
author = {McFie, Pamela J. and Stone, Sandra L. and Banman, Shanna L. and Stone, Scot J.},
doi = {10.1074/jbc.M110.163691},
issn = {00219258},
journal = {Journal of Biological Chemistry},
month = {nov},
number = {48},
pages = {37377--37387},
pmid = {20876538},
publisher = {Elsevier},
title = {{Topological orientation of acyl-CoA:Diacylglycerol acyltransferase-1 (DGAT1) and identification of a putative active site histidine and the role of the N terminus in dimer/tetramer formation}},
volume = {285},
year = {2010}
}
@article{Ward2021,
abstract = {Design of new drug compounds with target properties is a key area of research in generative modeling. We present a small drug molecule design pipeline based on graph-generative models and a comparison study of two state-of-the-art graph generative models for designing COVID-19 targeted drug candidates: 1) a variational autoencoder-based approach (VAE) that uses prior knowledge of molecules that have been shown to be effective for earlier coronavirus treatments and 2) a deep Q-learning method (DQN) that generates optimized molecules without any proximity constraints. We evaluate the novelty of the automated molecule generation approaches by validating the candidate molecules with drug-protein binding affinity models. The VAE method produced two novel molecules with similar structures to the antiretroviral protease inhibitor Indinavir that show potential binding affinity for the SARS-CoV-2 protein target 3-chymotrypsin-like protease (3CL-protease).},
archivePrefix = {arXiv},
arxivId = {2102.04977},
author = {Ward, Logan and Bilbrey, Jenna A. and Choudhury, Sutanay and Kumar, Neeraj and Sivaraman, Ganesh},
eprint = {2102.04977},
month = {feb},
title = {{Benchmarking Deep Graph Generative Models for Optimizing New Drug Molecules for COVID-19}},
year = {2021}
}
@article{Berardini2015,
author = {Berardini, Tanya Z. and Reiser, Leonore and Li, Donghui and Mezheritsky, Yarik and Muller, Robert and Strait, Emily and Huala, Eva},
doi = {10.1002/dvg.22877},
issn = {1526954X},
journal = {genesis},
month = {aug},
number = {8},
pages = {474--485},
title = {{The arabidopsis information resource: Making and mining the “gold standard” annotated reference plant genome}},
volume = {53},
year = {2015}
}
@article{Gammons2016,
author = {Gammons, Melissa V. and Renko, Miha and Johnson, Christopher M. and Rutherford, Trevor J. and Bienz, Mariann},
doi = {10.1016/j.molcel.2016.08.026},
issn = {10972765},
journal = {Molecular Cell},
month = {oct},
number = {1},
pages = {92--104},
title = {{Wnt Signalosome Assembly by DEP Domain Swapping of Dishevelled}},
volume = {64},
year = {2016}
}
@article{Wang2015,
author = {Wang, Hui and Klein, Michael G and Zou, Hua and Lane, Weston and Snell, Gyorgy and Levin, Irena and Li, Ke and Sang, Bi-Ching},
doi = {10.1038/nsmb.3049},
issn = {1545-9993},
journal = {Nature Structural {\&} Molecular Biology},
month = {jul},
number = {7},
pages = {581--585},
title = {{Crystal structure of human stearoyl–coenzyme A desaturase in complex with substrate}},
volume = {22},
year = {2015}
}
@article{Nachtschatt2020,
author = {Nachtschatt, Matthias and Okada, Shoko and Speight, Robert},
doi = {10.1002/ejlt.202000181},
issn = {1438-7697},
journal = {European Journal of Lipid Science and Technology},
month = {dec},
number = {12},
pages = {2000181},
title = {{Integral Membrane Fatty Acid Desaturases: A Review of Biochemical, Structural, and Biotechnological Advances}},
volume = {122},
year = {2020}
}
@incollection{HARWOOD1980,
title = {1 - Plant Acyl Lipids: Structure, Distribution, and Analysis},
editor = {P.K. Stumpf},
booktitle = {Lipids: Structure and Function},
publisher = {Academic Press},
pages = {1-55},
year = {1980},
isbn = {978-0-12-675404-9},
doi = {10.1016/B978-0-12-675404-9.50007-2},
author = {J.L. Harwood},
abstract = {Publisher Summary
Acyl lipids are major constituents of plant tissue. They comprise a wide variety of different structures of which the majority contain fatty acid esters, though in a few cases, the fatty acids are present in ether linkages or as amide derivatives. The lipids are mainly present as triacylglycerols in seeds or in the fleshy part of fruits where they act as food stores. These tissues provide important commercial sources of fats and oils. In other plant tissues such as leaves, roots, or shoots, acyl lipids are mainly present as glycolipids and phospholipids. The reason that acyl lipids are fats is because of their fatty acid constituents whose long hydrocarbon chains are basically hydrophobic. It is, therefore, logical to consider the fatty acids of plants first. Although the number of fatty acids detected in plant tissues approaches 300, most of them only occur in a few plant species. The major fatty acids are all saturated or unsaturated monocarboxylic acids with an unbranched even-numbered carbon chain.}
}
@article{Petrini2004,
author = {Petrini, Guillermo A. and Altabe, Silvia G. and Uttaro, Antonio D.},
doi = {10.1111/j.1432-1033.2004.04005.x},
issn = {00142956},
journal = {European Journal of Biochemistry},
month = {mar},
number = {6},
pages = {1079--1086},
title = {{Trypanosoma brucei oleate desaturase may use a cytochrome b5 -like domain in another desaturase as an electron donor}},
volume = {271},
year = {2004}
}
@article{Liu2012,
author = {Liu, Lin and Byeon, In-Ja L. and Bahar, Ivet and Gronenborn, Angela M.},
doi = {10.1021/ja210118w},
issn = {0002-7863},
journal = {Journal of the American Chemical Society},
month = {mar},
number = {9},
pages = {4229--4235},
title = {{Domain Swapping Proceeds via Complete Unfolding: A 19 F- and 1 H-NMR Study of the Cyanovirin-N Protein}},
volume = {134},
year = {2012}
}
@article{Pastor2015,
author = {Pastor, Nina and Amero, Carlos},
doi = {10.3389/fpls.2015.00306},
issn = {1664-462X},
journal = {Frontiers in Plant Science},
month = {may},
title = {{Information flow and protein dynamics: the interplay between nuclear magnetic resonance spectroscopy and molecular dynamics simulations}},
volume = {6},
year = {2015}
}
@article{Jasniewski2018,
author = {Jasniewski, Andrew J. and Que, Lawrence},
doi = {10.1021/acs.chemrev.7b00457},
issn = {0009-2665},
journal = {Chemical Reviews},
month = {mar},
number = {5},
pages = {2554--2592},
title = {{Dioxygen Activation by Nonheme Diiron Enzymes: Diverse Dioxygen Adducts, High-Valent Intermediates, and Related Model Complexes}},
volume = {118},
year = {2018}
}
@article{Haywood2018,
author = {Haywood, Elizabeth E. and Ho, Mengfei and Wilson, Brenda A.},
doi = {10.1074/jbc.RA117.001381},
issn = {00219258},
journal = {Journal of Biological Chemistry},
month = {mar},
number = {10},
pages = {3860--3870},
title = {{Modular domain swapping among the bacterial cytotoxic necrotizing factor (CNF) family for efficient cargo delivery into mammalian cells}},
volume = {293},
year = {2018}
}
@article{Consortium2019,
abstract = {The UniProt Knowledgebase is a collection of sequences and annotations for over 120 million proteins across all branches of life. Detailed annotations extracted from the literature by expert curators have been collected for over half a million of these proteins. These annotations are supplemented by annotations provided by rule based automated systems, and those imported from other resources. In this article we describe significant updates that we have made over the last 2 years to the resource. We have greatly expanded the number of Reference Proteomes that we provide and in particular we have focussed on improving the number of viral Reference Proteomes. The UniProt website has been augmented with new data visualizations for the subcellular localization of proteins as well as their structure and interactions. UniProt resources are available under a CC-BY (4.0) license via the web at https://www.uniprot.org/.},
author = {Consortium, The UniProt},
doi = {10.1093/nar/gky1049},
issn = {0305-1048},
journal = {Nucleic Acids Research},
month = {jan},
number = {D1},
pages = {D506--D515},
title = {{UniProt: a worldwide hub of protein knowledge}},
volume = {47},
year = {2019}
}
@article{Domergue2003,
abstract = {The health benefits attributed to very long-chain polyunsaturated fatty acids and the long term goal to produce them in transgenic oilseed crops have led to the cloning of all the genes coding for the desaturases and elongases involved in their biosynthesis. The encoded activities have been confirmed in vivo by heterologous expression, but very little is known about the actual acyl substrates involved in these pathways. Using a $\Delta$6-elongase and front-end desaturases from different organisms, we have reconstituted in Saccharomyces cerevisiae the biosynthesis of arachidonic acid from exogenously supplied linoleic acid in order to identify these acyl carriers. Acyl-CoA measurements strongly suggest that the elongation step involved in polyunsaturated fatty acids biosynthesis is taking place within the acyl-CoA pool. In contrast, detailed analyses of lipids revealed that the two desaturation steps ($\Delta$5 and $\Delta$6) occur predominantly at the sn-2 position of phosphatidylcholine when using $\Delta$5- and $\Delta$6-desaturases from lower plants, fungi, worms, and algae. The specificity of these $\Delta$6-desaturases for the fatty acid acylated at this particular position as well as a limiting re-equilibration with the acyl-CoA pool result in the accumulation of $\gamma$-linolenic acid at the sn-2 position of phosphatidylcholine and prevent efficient arachidonic acid biosynthesis in yeast. We confirm by using a similar experimental approach that, in contrast, the human $\Delta$6-desaturase uses linoleoyl-CoA as substrate, which results in high efficiency of the subsequent elongation step. In addition, we report that $\Delta$12-desaturases have no specificity toward the lipid polar headgroup or the sn-position.},
author = {Domergue, Fr{\'{e}}d{\'{e}}ric and Abbadi, Amine and Ott, Claudia and Zank, Thorsten K. and Z{\"{a}}hringer, Ulrich and Heinz, Ernst},
doi = {10.1074/jbc.M305990200},
issn = {00219258},
journal = {Journal of Biological Chemistry},
month = {sep},
number = {37},
pages = {35115--35126},
pmid = {12835316},
publisher = {Elsevier},
title = {{Acyl carriers used as substrates by the desaturases and elongases involved in very long-chain polyunsaturated fatty acids biosynthesis reconstituted in yeast}},
volume = {278},
year = {2003}
}
@misc{Pronk1996,
abstract = {In yeasts, pyruvate is located at a major junction of assimilatory and dissimilatory reactions as well as at the branch-point between respiratory dissimilation of sugars and alcoholic fermentation. This review deals with the enzymology, physiological function and regulation of three key reactions occurring at the pyruvate branch-point in the yeast Saccharomyces cerevisiae (i) the direct oxidative decarboxylation of pyruvate to acetyl-CoA, catalysed by the pyruvate dehydrogenase complex, (ii) decarboxylation of pyruvate to acetaldehyde, catalysed by pyruvate decarboxylase, and (iii) the anaplerotic carboxylation of pyruvate to oxaloacetate, catalysed by pyruvate carboxylase. Special attention is devoted to physiological studies on S. cerevisiae strains in which structural genes encoding these key enzymes have been inactivated by gene disruption.},
author = {Pronk, Jack T. and Steensma, H. Yde and {Van Dijken}, Johannes P.},
booktitle = {Yeast},
doi = {10.1002/(SICI)1097-0061(199612)12:16<1607::AID-YEA70>3.0.CO;2-4},
issn = {0749503X},
keywords = {Alcoholic fermentation,Crabtree effect,Glycolysis,Metabolic engineering,Pyruvate carboxylase,Pyruvate decaroxylase,Pyruvate dehydrogenase complex,Sugar metabolism,TCA cycle,Yeast},
month = {dec},
number = {16},
pages = {1607--1633},
pmid = {9123965},
title = {{Pyruvate metabolism in Saccharomyces cerevisiae}},
volume = {12},
year = {1996}
}
@article{Okuley1994,
abstract = {The polyunsaturated fatty acids linoleate and alpha-linolenate are important membrane components and are the essential fatty acids of human nutrition. The major enzyme responsible for the synthesis of these compounds is the plant oleate desaturase of the endoplasmic reticulum, and its activity is controlled in Arabidopsis by the fatty acid desaturation 2 (fad2) locus. A fad2 allele was identified in a population of Arabidopsis in which mutations had been created by T-DNA insertions. Genomic DNA flanking the T-DNA was cloned by plasmid rescue and used to isolate cDNA and genomic clones of FAD2. A cDNA containing the entire FAD2 coding sequence was expressed in fad2 mutant plants and shown to complement the mutant fatty acid phenotype. The deduced amino acid sequence from the cDNA showed homology to other plant desaturases, and this confirmed that FAD2 is the structural gene for the desaturase. Gel blot analyses of FAD2 mRNA levels showed that the gene is expressed throughout the plant and suggest that transcript levels are in excess of the amount needed to account for oleate desaturation. Sequence analysis identified histidine-rich motifs that could contribute to an iron binding site in the cytoplasmic domain of the protein. Such a position would facilitate interaction between the desaturase and cytochrome b5, which is the direct source of electrons for the desaturation reaction, but would limit interaction of the active site with the fatty acyl substrate.},
author = {Okuley, J and Lightner, J and Feldmann, K and Yadav, N and Lark, E and Browse, J},
doi = {10.1105/tpc.6.1.147},
issn = {1040-4651},
journal = {The Plant cell},
keywords = {Alleles,Amino Acid Sequence,Arabidopsis/enzymology/*genetics,Base Sequence,Cloning, Molecular,Cold Temperature,DNA,DNA, Bacterial,Fatty Acid Desaturases/*genetics/metabolism,Genes, Plant,Genetic Complementation Test,Iron/metabolism,Linoleic Acid,Linoleic Acids/*biosynthesis,Molecular Sequence Data,Mutation,Oxidoreductases Acting on CH-CH Group Donors,Plant Proteins/genetics/metabolism,Sequence Homology, Amino Acid,Transcription, Genetic,alpha-Linolenic Acid/*biosynthesis},
language = {eng},
month = {jan},
number = {1},
pages = {147--158},
title = {{Arabidopsis FAD2 gene encodes the enzyme that is essential for polyunsaturated lipid synthesis}},
volume = {6},
year = {1994}
}
@article{Lou2014,
author = {Lou, Ying and Schwender, Jorg and Shanklin, John},
doi = {10.1074/jbc.M114.572883},
issn = {00219258},
journal = {Journal of Biological Chemistry},
month = {jun},
number = {26},
pages = {17996--18007},
title = {{FAD2 and FAD3 Desaturases Form Heterodimers That Facilitate Metabolic Channeling in Vivo}},
volume = {289},
year = {2014}
}
@article{Liu2008,
author = {Liu, J. and Nussinov, R.},
doi = {10.1073/pnas.0707401105},
issn = {0027-8424},
journal = {Proceedings of the National Academy of Sciences},
month = {jan},
number = {3},
pages = {901--906},
title = {{Allosteric effects in the marginally stable von Hippel-Lindau tumor suppressor protein and allostery-based rescue mutant design}},
volume = {105},
year = {2008}
}
@article{Covello1996,
author = {Covello, P. S. and Reed, D. W.},
doi = {10.1104/pp.111.1.223},
issn = {0032-0889},
journal = {Plant Physiology},
month = {may},
number = {1},
pages = {223--226},
title = {{Functional Expression of the Extraplastidial Arabidopsis thaliana Oleate Desaturase Gene (FAD2) in Saccharomyces cerevisiae}},
volume = {111},
year = {1996}
}
@article{Kelley2015,
abstract = {Phyre2 is a suite of tools available on the web to predict and analyze protein structure, function and mutations. The focus of Phyre2 is to provide biologists with a simple and intuitive interface to state-of-the-art protein bioinformatics tools. Phyre2 replaces Phyre, the original version of the server for which we previously published a paper in Nature Protocols. In this updated protocol, we describe Phyre2, which uses advanced remote homology detection methods to build 3D models, predict ligand binding sites and analyze the effect of amino acid variants (e.g., nonsynonymous SNPs (nsSNPs)) for a user's protein sequence. Users are guided through results by a simple interface at a level of detail they determine. This protocol will guide users from submitting a protein sequence to interpreting the secondary and tertiary structure of their models, their domain composition and model quality. A range of additional available tools is described to find a protein structure in a genome, to submit large number of sequences at once and to automatically run weekly searches for proteins that are difficult to model. The server is available at http://www.sbg.bio.ic.ac.uk/phyre2. A typical structure prediction will be returned between 30 min and 2 h after submission.},
author = {Kelley, Lawrence A. and Mezulis, Stefans and Yates, Christopher M. and Wass, Mark N. and Sternberg, Michael J.E.},
doi = {10.1038/nprot.2015.053},
issn = {17502799},
journal = {Nature Protocols},
keywords = {Bioinformatics,Protein function predictions,Protein structure predictions,Structural biology},
month = {jun},
number = {6},
pages = {845--858},
pmid = {25950237},
publisher = {Nature Publishing Group},
title = {{The Phyre2 web portal for protein modeling, prediction and analysis}},
volume = {10},
year = {2015}
}
@article{Sun2017,
author = {Sun, Zhaoyang and {El Omari}, Kamel and Sun, Xiaoyu and Ilca, Serban L. and Kotecha, Abhay and Stuart, David I. and Poranen, Minna M. and Huiskonen, Juha T.},
doi = {10.1038/ncomms14814},
issn = {2041-1723},
journal = {Nature Communications},
month = {apr},
number = {1},
pages = {14814},
title = {{Double-stranded RNA virus outer shell assembly by bona fide domain-swapping}},
volume = {8},
year = {2017}
}
@article{Alloatti2011,
abstract = {A detailed analysis of the trypanosomatids' genome projects revealed the presence of genes predicted to encode fatty-acid desaturases of the methyl-end type (MED). After cloning and functional characterization of all identified genes, it can be concluded that Trypanosoma cruzi contains two MEDs with oleate desaturase (OD) activities whereas Leishmania major contains one OD and two active linoleate desaturases (LD). All characterized ODs are highly specific for oleate (18:1$\Delta$9) as substrate, presenting a $\nu$ + 3 regioselectivity, although palmitoleate (16:1$\Delta$9) can be desaturated as well, but to a lesser extent. L. major LD appears to use exclusively linoleate (18:2n-6), converting it into $\alpha$-linolenate (18:3n-3). This strong specificity assures no further conversion of polyunsaturated fatty acids (PUFAs) of the n-6 series into the n-3 series, downstream in the PUFA biosynthesis pathway. This characterization completes the identification of all enzymes involved in PUFA biosynthesis in a parasitic protist. Differently from their Trypanosoma brucei orthologue, T. cruzi and L. major ODs were more active when expressed either, in the presence of trienoic fatty acids or at higher temperatures. This could be evidence for a differential post-translational regulation of these enzymes as a result of direct sensing of environmentally dependent parameters such as membrane fluidity. {\textcopyright} 2010 Elsevier B.V. All rights reserved.},
author = {Alloatti, Andr{\'{e}}s and Uttaro, Antonio D.},
doi = {10.1016/j.molbiopara.2010.10.006},
issn = {01666851},
journal = {Molecular and Biochemical Parasitology},
keywords = {Leishmania,Methyl-end desaturases,Regioselectivity,Trypanosoma,Unsaturated fatty acids},
month = {feb},
number = {2},
pages = {126--132},
pmid = {20974196},
publisher = {Elsevier},
title = {{Highly specific methyl-end fatty-acid desaturases of trypanosomatids}},
volume = {175},
year = {2011}
}
@article{Petrie2012,
abstract = {Background: Monoacylglycerol acyltransferases (MGATs) are predominantly associated with lipid absorption and resynthesis in the animal intestine where they catalyse the first step in the monoacylglycerol (MAG) pathway by acylating MAG to form diacylglycerol (DAG). Typical plant triacylglycerol (TAG) biosynthesis routes such as the Kennedy pathway do not include an MGAT step. Rather, DAG and TAG are synthesised de novo from glycerol-3-phosphate (G-3-P) by a series of three subsequent acylation reactions although a complex interplay with membrane lipids exists. Methodology/Principal Findings: We demonstrate that heterologous expression of a mouse MGAT acyltransferase in Nicotiana benthamiana significantly increases TAG accumulation in vegetative tissues despite the low levels of endogenous MAG substrate available. In addition, DAG produced by this acyltransferase can serve as a substrate for both native and coexpressed diacylglycerol acyltransferases (DGAT). Finally, we show that the Arabidopsis thaliana GPAT4 acyltransferase can produce MAG in Saccharomyces cerevisiae using oleoyl-CoA as the acyl-donor. Conclusions/Significance: This study demonstrates the concept of a new method of increasing oil content in vegetative tissues by using MAG as a substrate for TAG biosynthesis. Based on in vitro yeast assays and expression results in N. benthamiana, we propose that co-expression of a MAG synthesising enzyme such as A. thaliana GPAT4 and a MGAT or bifunctional M/DGAT can result in DAG and TAG synthesis from G-3-P via a route that is independent and complementary to the endogenous Kennedy pathway and other TAG synthesis routes. {\textcopyright} 2012 Petrie et al.},
author = {Petrie, James R. and Vanhercke, Thomas and Shrestha, Pushkar and {El Tahchy}, Anna and White, Adam and Zhou, Xue-Rong and Liu, Qing and Mansour, Maged P. and Nichols, Peter D. and Singh, Surinder P.},
doi = {10.1371/annotation/f0547e8e-5cec-4ae4-8348-07f5aa3b8f34},
issn = {1932-6203},
journal = {PLoS ONE},
keywords = {Arabidopsis thaliana,Fatty acids,Genetically modified plants,Leaves,Lipids,Neutral lipids,Saccharomyces cerevisiae,Yeast},
month = {jul},
number = {7},
pages = {e35214},
publisher = {Public Library of Science (PLoS)},
title = {{Correction: Recruiting a New Substrate for Triacylglycerol Synthesis in Plants: The Monoacylglycerol Acyltransferase Pathway}},
volume = {7},
year = {2012}
}
@article{Gagne2009,
author = {Gagn\'e, Steve J. and Reed, Darwin W. and Gray, Gordon R. and Covello, Patrick S.},
doi = {10.1021/bi901605d},
issn = {0006-2960},
journal = {Biochemistry},
month = {dec},
number = {51},
pages = {12298--12304},
title = {{Structural Control of Chemoselectivity, Stereoselectivity, and Substrate Specificity in Membrane-Bound Fatty Acid Acetylenases and Desaturases}},
volume = {48},
year = {2009}
}
@article{Park2010,
author = {Park, Chad K. and Joshi, Hemant K. and Agrawal, Alka and Ghare, M. Imran and Little, Elizabeth J. and Dunten, Pete W. and Bitinaite, Jurate and Horton, Nancy C.},
doi = {10.1371/journal.pbio.1000554},
editor = {Petsko, Gregory A.},
issn = {1545-7885},
journal = {PLoS Biology},
month = {dec},
number = {12},
pages = {e1000554},
title = {{Domain Swapping in Allosteric Modulation of DNA Specificity}},
volume = {8},
year = {2010}
}
@article{Meesapyodsuk2014,
author = {Meesapyodsuk, Dauenpen and Qiu, Xiao},
doi = {10.1021/cb400675d},
issn = {1554-8929},
journal = {ACS Chemical Biology},
month = {apr},
number = {4},
pages = {922--934},
title = {{Structure Determinants for the Substrate Specificity of Acyl-CoA {$\Delta$}9 Desaturases from a Marine Copepod}},
volume = {9},
year = {2014}
}
@article{Singh2001,
abstract = {The Crepis palaestina cDNA Cpal2 encodes a $\Delta$12-epoxygenase that can catalyse the synthesis of 12,13-epoxy-cis-9-octadecenoic acid (18:1E) from linoleic acid (18:2). When the Cpal2 gene was expressed under the control of the napin seed-specific promoter in Arabidopsis thaliana (L.) Heynh., the seed lipids accumulated only low levels of 18:1E and also 12,13-epoxy-cis-9,15-octadec-2-enoic acid (18:2E). Despite the fact that the levels of these epoxy fatty acids comprised only up to 6.2{\%} of the total fatty acids, there was a very marked increase in oleic acid (18:1) and decrease in linoleic (18:2) and $\alpha$-linolenic (18:3) acids in these plants, indicating that endogenous $\Delta$12-desaturation was greatly reduced in these plants. Significant between-line differences in the levels of Cpal2 mRNA were observed during seed development, but were not associated with any major variation in mRNA levels for the endogenous Arabidopsis$\Delta$12-desaturase (Fad2). This suggests that if an unfavourable interaction occurs between the transgenic $\Delta$12-epoxygenase and the endogenous $\Delta$12-desaturase, which decreases the level of desaturation, it occurs at either the translational or post-translational level. We further show that the co-expression of a $\Delta$12-desaturase gene from C. palaestina in Cpal2 transgenic Arabidopsis returns the relative proportions of the C18 seed fatty acids to normal levels and results in an almost twofold increase in total epoxy fatty acids.},
author = {Singh, Surinder and Thomaeus, Stefan and Lee, Michael and Stymne, Sten and Green, Allan},
doi = {10.1007/s004250000456},
issn = {1432-2048},
journal = {Planta},
number = {5},
pages = {872--879},
title = {{Transgenic expression of a $\Delta$12-epoxygenase gene in Arabidopsis seeds inhibits accumulation of linoleic acid}},
volume = {212},
year = {2001}
}
@article{Lua2016,
author = {Lua, Rhonald C. and Wilson, Stephen J. and Konecki, Daniel M. and Wilkins, Angela D. and Venner, Eric and Morgan, Daniel H. and Lichtarge, Olivier},
doi = {10.1093/nar/gkv1279},
issn = {0305-1048},
journal = {Nucleic Acids Research},
month = {jan},
number = {D1},
pages = {D308--D312},
title = {{UET: a database of evolutionarily-predicted functional determinants of protein sequences that cluster as functional sites in protein structures}},
volume = {44},
year = {2016}
}
@article{Shi2020,
author = {Shi, Haisu and Tian, Jinlong and Wu, Chen and Li, Mo and An, Feiyu and Wu, Rina and Shao, Junhua and Zheng, Yan and Luo, Xue and Tao, Dongbing and Chen, Xu and Pi, Yuzhen and Zhao, Chunyan and Yue, Xiqing and Wu, Junrui},
doi = {10.1016/j.enzmictec.2020.109563},
issn = {01410229},
journal = {Enzyme and Microbial Technology},
month = {aug},
pages = {109563},
title = {{Determination of allosteric and active sites responsible for catalytic activity of delta 12 fatty acid desaturase from Geotrichum candidum and Mortierella alpina by domain swapping}},
volume = {138},
year = {2020}
}
@article{Berman2000,
abstract = {The Protein Data Bank (PDB; http://www.rcsb.org/pdb/ ) is the single worldwide archive of structural data of biological macromolecules. This paper describes the goals of the PDB, the systems in place for data deposition and access, how to obtain further information, and near-term plans for the future development of the resource.},
author = {Berman, Helen M and Westbrook, John and Feng, Zukang and Gilliland, Gary and Bhat, T N and Weissig, Helge and Shindyalov, Ilya N and Bourne, Philip E},
doi = {10.1093/nar/28.1.235},
issn = {0305-1048},
journal = {Nucleic Acids Research},
month = {jan},
number = {1},
pages = {235--242},
title = {{The Protein Data Bank}},
volume = {28},
year = {2000}
}
@article{Mekhedov2000,
author = {Mekhedov, Sergei and de Il{\'{a}}rduya, Oskar Martı́nez and Ohlrogge, John},
doi = {10.1104/pp.122.2.389},
issn = {0032-0889},
journal = {Plant Physiology},
month = {feb},
number = {2},
pages = {389--402},
title = {{Toward a Functional Catalog of the Plant Genome. A Survey of Genes for Lipid Biosynthesis}},
volume = {122},
year = {2000}
}
@article{Qin2016,
author = {Qin, Bin and Matsuda, Yudai and Mori, Takahiro and Okada, Masahiro and Quan, Zhiyang and Mitsuhashi, Takaaki and Wakimoto, Toshiyuki and Abe, Ikuro},
doi = {10.1002/anie.201509263},
issn = {14337851},
journal = {Angewandte Chemie International Edition},
month = {jan},
number = {5},
pages = {1658--1661},
title = {{An Unusual Chimeric Diterpene Synthase from Emericella variecolor and Its Functional Conversion into a Sesterterpene Synthase by Domain Swapping}},
volume = {55},
year = {2016}
}
@article{Cao2003,
abstract = {Acyl-CoA:monoacylglycerol acyltransferase (MGAT) plays an important role in dietary fat absorption by catalyzing a rate-limiting step in the re-synthesis of diacylglycerols in enterocytes. The present study reports further characterization of MGAT2, a newly identified intestinal MGAT (Cao, J., Lockwood, J., Burn, P., and Shi, Y. (2003) J. Biol. Chem. 278, 13860-13866) for its substrate specificity, requirement for lipid cofactors, optimum pH and Mg2+, and other intrinsic properties. MGAT2 enzyme expressed in COS-7 cells displayed a broad range of substrate specificity toward fatty acyl-CoA derivatives and monoacylglycerols, among which the highest activities were observed with oleoyl-CoA and rac-1-monolauroylglycerol, respectively. MGAT2 appeared to acylate monoacylglycerols containing unsaturated fatty acyls in preference to saturated ones. Lipid cofactors that play roles in signal transduction were shown to modulate MGAT2 activities. In contrast to oleic acid and sphingosine that exhibited inhibitory effects, phosphatidylcholine, phosphatidylserine, and phosphatidic acid stimulated MGAT2 activities. Using recombinant murine MGAT2 expressed in Escherichia coli, we demonstrated conclusively that MGAT2 also possessed an intrinsic acyl-CoA:diacylglycerol acyltransferase (DGAT) activity, which could provide an alternative pathway for triacylglycerol synthesis in the absence of DGAT. In contrast to the inhibitory effect on MGAT2 activities, nonionic and zwitterionic detergents led to a striking activation of DGAT activity of the human DGAT1 expressed in mammalian cells, which further distinguished the behaviors of the two enzymes. The elucidation of properties of MGAT2 will facilitate future development of compounds that inhibit dietary fat absorption as a means to treat obesity.},
author = {Cao, Jingsong and Burn, Paul and Shi, Yuguang},
doi = {10.1074/jbc.M302835200},
issn = {00219258},
journal = {Journal of Biological Chemistry},
month = {jul},
number = {28},
pages = {25657--25663},
pmid = {12730219},
publisher = {Elsevier},
title = {{Properties of the mouse intestinal Acyl-CoA:monoacylglycerol acyltransferase, MGAT2}},
volume = {278},
year = {2003}
}
@article{Hoffmann2008,
abstract = {$\omega$3-Very long chain polyunsaturated fatty acids (VLCPUFA) are essential for human development and brain function and, thus, are indispensable components of the human diet. The current main source of VLCPUFA sis represented by ocean fish stocks, which are in severe decline, and the development of alternative, sustainable sources of VLCPUFAs is urgently required. Our research aims at exploiting the powerful infrastructure available for the large scale culture of oilseed crops, such as rapeseed, to produce VLCPUFAs such as eicosapentaenoic acid in transgenic plants. VLCPUF Abiosynthesis requires repeated desaturation and repeated elongation of long chain fatty acid substrates. In previous experiments the production of eicosapentaenoic acid in transgenic plants was found to be limited by an unexpected bottleneck represented by the acyl exchange between the site of desaturation, endoplasmic reticulum-associated phospholipids, and the site of elongation, the cytosolic acyl-CoA pool. Here we report on the establishment of a coordinated, exclusively acyl-CoA-dependent pathway, which avoids the rate-limiting transesterification steps between the acyl lipids and the acyl-CoA pool during VLCPUFA biosynthesis. The pathway is defined by previously uncharacterized enzymes, encoded by cDNAs isolated from the microalga Mantoniella squamata. The conceptual enzymatic pathway was established and characterized first in yeast to provide proof-ofconcept data for its feasibility and subsequently in seeds of Arabidopsis thaliana. The comparison of the acyl-CoA-dependent pathway with the known lipid-linked pathway for VLCPUFA biosynthesis showed that the acyl-CoA-dependent pathway circumvents the bottleneck of switching the $\Delta$6-desaturated fatty acids between lipids and acyl-CoA in Arabidopsis seeds. {\textcopyright} 2008 by The American Society for Biochemistry and Molecular Biology, Inc.},
author = {Hoffmann, Mareike and Wagner, Martin and Abbadi, Amine and Fulda, Martin and Feussner, Ivo},
doi = {10.1074/jbc.M802377200},
issn = {00219258},
journal = {Journal of Biological Chemistry},
month = {aug},
number = {33},
pages = {22352--22362},
pmid = {18556660},
publisher = {Elsevier},
title = {{Metabolic engineering of $\omega$3-very long chain polyunsaturated fatty acid production by an exclusively acyl-CoA-dependent pathway}},
volume = {283},
year = {2008}
}
@article{Behrouzian2001,
author = {Behrouzian, Behnaz and Fauconnot, La{\"{e}}titia and Daligault, Franck and Nugier-Chauvin, Caroline and Patin, Henri and Buist, Peter H.},
doi = {10.1046/j.1432-1327.2001.02258.x},
issn = {00142956},
journal = {European Journal of Biochemistry},
month = {jun},
number = {12},
pages = {3545--3549},
title = {{Mechanism of fatty acid desaturation in the green alga Chlorella vulgaris}},
volume = {268},
year = {2001}
}
@article{Fox1993,
author = {Fox, B. G. and Shanklin, J. and Somerville, C. and Munck, E.},
doi = {10.1073/pnas.90.6.2486},
issn = {0027-8424},
journal = {Proceedings of the National Academy of Sciences},
month = {mar},
number = {6},
pages = {2486--2490},
title = {{Stearoyl-acyl carrier protein delta 9 desaturase from Ricinus communis is a diiron-oxo protein.}},
volume = {90},
year = {1993}
}
@article{Vettoretti2016,
author = {Vettoretti, Gerolamo and Moroni, Elisabetta and Sattin, Sara and Tao, Jiahui and Agard, David A. and Bernardi, Anna and Colombo, Giorgio},
doi = {10.1038/srep23830},
issn = {2045-2322},
journal = {Scientific Reports},
month = {jul},
number = {1},
pages = {23830},
title = {{Molecular Dynamics Simulations Reveal the Mechanisms of Allosteric Activation of Hsp90 by Designed Ligands}},
volume = {6},
year = {2016}
}
@article{Zhang2008,
author = {Zhang, Yong-Mei and Rock, Charles O.},
doi = {10.1038/nrmicro1839},
issn = {1740-1526},
journal = {Nature Reviews Microbiology},
month = {mar},
number = {3},
pages = {222--233},
title = {{Membrane lipid homeostasis in bacteria}},
volume = {6},
year = {2008}
}
@article{Gupta1977,
author = {Gupta, C. M. and Radhakrishnan, R. and Khorana, H. G.},
doi = {10.1073/pnas.74.10.4315},
issn = {0027-8424},
journal = {Proceedings of the National Academy of Sciences},
month = {oct},
number = {10},
pages = {4315--4319},
title = {{Glycerophospholipid synthesis: improved general method and new analogs containing photoactivable groups.}},
volume = {74},
year = {1977}
}
@article{Whittle2020,
author = {Whittle, Edward J. and Cai, Yuanheng and Keereetaweep, Jantana and Chai, Jin and Buist, Peter H. and Shanklin, John},
doi = {10.1104/pp.19.01111},
issn = {0032-0889},
journal = {Plant Physiology},
month = {feb},
number = {2},
pages = {730--738},
title = {{Castor Stearoyl-ACP Desaturase Can Synthesize a Vicinal Diol by Dioxygenase Chemistry}},
volume = {182},
year = {2020}
}
@article{Vendome2011,
author = {Vendome, Jeremie and Posy, Shoshana and Jin, Xiangshu and Bahna, Fabiana and Ahlsen, Goran and Shapiro, Lawrence and Honig, Barry},
doi = {10.1038/nsmb.2051},
issn = {1545-9993},
journal = {Nature Structural {\&} Molecular Biology},
month = {jun},
number = {6},
pages = {693--700},
title = {{Molecular design principles underlying $\beta$-strand swapping in the adhesive dimerization of cadherins}},
volume = {18},
year = {2011}
}
@article{Hassan2017,
abstract = {"Virtual Screening" is a common step of in silico drug design, where researchers screen a large library of small molecules (ligands) for interesting hits, in a process known as "Docking". However, docking is a computationally intensive and time-consuming process, usually restricted to small size binding sites (pockets) and small number of interacting residues. When the target site is not known (blind docking), researchers split the docking box into multiple boxes, or repeat the search several times using different seeds, and then merge the results manually. Otherwise, the search time becomes impractically long. In this research, we studied the relation between the search progression and Average Sum of Proximity relative Frequencies (ASoF) of searching threads, which is closely related to the search speed and accuracy. A new inter-process spatio-temporal integration method is employed in Quick Vina 2, resulting in a new docking tool, QuickVina-W, a suitable tool for "blind docking", (not limited in search space size or number of residues). QuickVina-W is faster than Quick Vina 2, yet better than AutoDock Vina. It should allow researchers to screen huge ligand libraries virtually, in practically short time and with high accuracy without the need to define a target pocket beforehand.},
author = {Hassan, Nafisa M. and Alhossary, Amr A. and Mu, Yuguang and Kwoh, Chee Keong},
doi = {10.1038/s41598-017-15571-7},
issn = {20452322},
journal = {Scientific Reports},
keywords = {Software,Virtual drug screening,Virtual screening},
month = {dec},
number = {1},
pages = {1--13},
pmid = {29133831},
publisher = {Nature Publishing Group},
title = {{Protein-Ligand Blind Docking Using QuickVina-W with Inter-Process Spatio-Temporal Integration}},
volume = {7},
year = {2017}
}
@article{Petrie2010,
abstract = {Long-chain (≥C20) polyunsaturated fatty acids (LC-PUFA) EPA and DHA (20:5$\Delta$5,8,11,14,17 and 22:6$\Delta$4,7,10,13,16,19) have well-documented health benefits against coronary heart disease, rheumatoid arthritis and other disorders. Currently, the predominant sources of these fatty acids are marine fish and algal oils, but research is being conducted to ensure that a sustainable, land-based production system can be developed. We here describe the metabolic engineering of an artificial pathway that produces 26{\%} EPA in leaf triacylglycerol using a newly-identified $\Delta$6-desaturase from the marine microalga Micromonas pusilla. We also demonstrate that this enzyme appears to function as an acyl-CoA desaturase that has preference for $\omega$3 substrates both in planta and in yeast. Phylogenetic analysis indicates that this desaturase shares highly conserved motifs with previously described acyl-CoA $\Delta$6-desaturases. Crown Copyright {\textcopyright} 2009.},
author = {Petrie, James R. and Shrestha, Pushkar and Mansour, Maged P. and Nichols, Peter D. and Liu, Qing and Singh, Surinder P.},
doi = {10.1016/j.ymben.2009.12.001},
issn = {10967176},
journal = {Metabolic Engineering},
keywords = {Acyl-CoA,LC-PUFA,Micromonas pusilla,Omega-3,SDA,$\Delta$6-desaturase},
month = {may},
number = {3},
pages = {233--240},
pmid = {20004733},
publisher = {Academic Press},
title = {{Metabolic engineering of omega-3 long-chain polyunsaturated fatty acids in plants using an acyl-CoA {$\Delta$}6-desaturase with $\omega$3-preference from the marine microalga Micromonas pusilla}},
volume = {12},
year = {2010}
}
@article{STUMPF1980,
author = {Stumpf, P.K.},
doi = {10.1016/B978-0-12-675404-9.50013-8},
journal = {Lipids: Structure and Function},
month = {jan},
pages = {177--204},
publisher = {Academic Press},
title = {{Biosynthesis of Saturated and Unsaturated Fatty Acids}},
year = {1980}
}
@article{Los1998,
abstract = {Fatty acid desaturases are enzymes that introduce double bonds into fatty acyl chains. They are present in all groups of organisms, i.e., bacteria, fungi, plants and animals, and play a key role in the maintenance of the proper structure and functioning of biological membranes. The desaturases are characterized by the presence of three conserved histidine tracks which are presumed to compose the Fe-binding active centers of the enzymes. Recent findings on the structure and expression of different types of fatty acid desaturase in cyanobacteria, plants and animals are reviewed in this article. Roles of individual desaturases in temperature acclimation and principles of regulation of the desaturase genes are discussed. Copyright (C) 1998.},
author = {Los, Dmitry A. and Murata, Norio},
doi = {10.1016/S0005-2760(98)00091-5},
issn = {0005-2760},
journal = {Biochimica et Biophysica Acta (BBA) - Lipids and Lipid Metabolism},
keywords = {Acclimation,Fatty acid desaturase,Gene expression,Gene regulation,Lipid,Membrane},
month = {oct},
number = {1},
pages = {3--15},
publisher = {Elsevier},
title = {{Structure and expression of fatty acid desaturases}},
volume = {1394},
year = {1998}
}
@article{Stubbs1984,
author = {Stubbs, Christopher D. and Smith, Anthony D.},
doi = {10.1016/0304-4157(84)90005-4},
issn = {0304-4157},
journal = {Biochimica et Biophysica Acta (BBA) - Reviews on Biomembranes},
month = {jan},
number = {1},
pages = {89--137},
publisher = {Elsevier},
title = {{The modification of mammalian membrane polyunsaturated fatty acid composition in relation to membrane fluidity and function}},
volume = {779},
year = {1984}
}
@article{Liu2014,
author = {Liu, Hui and Cheng, Tao and Xian, Mo and Cao, Yujin and Fang, Fang and Zou, Huibin},
doi = {10.1016/j.biotechadv.2013.12.003},
issn = {07349750},
journal = {Biotechnology Advances},
month = {mar},
number = {2},
pages = {382--389},
title = {{Fatty acid from the renewable sources: A promising feedstock for the production of biofuels and biobased chemicals}},
volume = {32},
year = {2014}
}
@article{Zhou2014,
abstract = {Volatile energy costs and environmental concerns have spurred interest in the development of alternative, renewable, sustainable and cost-effective energy resources. Advanced biofuels have potential to replace fossil fuels in supporting high-power demanding machinery such as aircrafts and trucks. Microbial biosynthesis is generally considered as an environmental friendly refinery process, and fatty acid biosynthesis is an attractive route to synthesize chemicals and especially drop-in biofuels due to the high degree of reduction of fatty acids. The robustness and excellent accessibility to molecular genetics make the yeast S. cerevisiae a suitable host for the production of biofuels, chemicals and pharmaceuticals, and recent advances in metabolic engineering as well as systems and synthetic biology allow us to engineer the yeast fatty acid metabolism and modification pathways for production of advanced biofuels and chemicals.},
author = {Zhou, Yongjin J. and Buijs, Nicolaas A. and Siewers, Verena and Nielsen, Jens},
doi = {10.3389/FBIOE.2014.00032},
issn = {2296-4185},
journal = {Frontiers in Bioengineering and Biotechnology},
keywords = {alkanes/alkene,fatty acid ethyl ester,fatty acid metabolism,fatty alcohol,yeast},
number = {SEP},
pages = {32},
publisher = {Frontiers},
title = {{Fatty Acid-Derived Biofuels and Chemicals Production in Saccharomyces cerevisiae}},
volume = {0},
year = {2014}
}
@article{Yu2014,
abstract = {Fatty acid derivatives, such as hydroxy fatty acids, fatty alcohols, fatty acid methyl/ethyl esters and fatty alka(e)nes, have a wide range of industrial applications including plastics, lubricants and fuels. Currently, these chemicals are obtained mainly through chemical synthesis, which is complex and costly, and their availability from natural biological sources is extremely limited. Metabolic engineering of microorganisms has provided a platform for effective production of these valuable biochemicals. Notably, synthetic biology-based metabolic engineering strategies have been extensively applied to refactor microorganisms for improved biochemical production. Here, we reviewed (i) the current status of metabolic engineering of microbes that produce fatty acid-derived valuable chemicals, and (ii) the recent progress of synthetic biology approaches that assist metabolic engineering, such as mRNA secondary structure engineering, sensor-regulator system, regulatable expression system, ultrasensitive input/output control system, and computer science-based design of complex gene circuits. Furthermore, key challenges and strategies were discussed. Finally, we concluded that synthetic biology provides useful metabolic engineering strategies for economically viable production of fatty acid-derived valuable chemicals in engineered microbes.},
author = {Yu, Ai-Qun and {Pratomo Juwono}, Nina Kurniasih and Leong, Susanna Su Jan and Chang, Matthew Wook},
doi = {10.3389/FBIOE.2014.00078},
issn = {2296-4185},
journal = {Frontiers in Bioengineering and Biotechnology},
keywords = {Escherichia coli,Metabolic Engineering,Synthetic Biology,biochemical production,fatty acid biosynthesis pathway,yeast},
number = {DEC},
pages = {78},
publisher = {Frontiers},
title = {{Production of Fatty Acid-Derived Valuable Chemicals in Synthetic Microbes}},
volume = {0},
year = {2014}
}
@article{Tee2014,
author = {Tee, Ting Wei and Chowdhury, Anupam and Maranas, Costas D. and Shanks, Jacqueline V.},
doi = {10.1002/bit.25205},
issn = {0006-3592},
journal = {Biotechnology and Bioengineering},
month = {may},
number = {5},
pages = {849--857},
title = {{Systems metabolic engineering design: Fatty acid production as an emerging case study}},
volume = {111},
year = {2014}
}
@article{Runguphan2014,
author = {Runguphan, Weerawat and Keasling, Jay D.},
doi = {10.1016/j.ymben.2013.07.003},
issn = {10967176},
journal = {Metabolic Engineering},
month = {jan},
pages = {103--113},
title = {{Metabolic engineering of Saccharomyces cerevisiae for production of fatty acid-derived biofuels and chemicals}},
volume = {21},
year = {2014}
}
@article{Lennen2013,
author = {Lennen, Rebecca M and Pfleger, Brian F},
doi = {10.1016/j.copbio.2013.02.028},
issn = {09581669},
journal = {Current Opinion in Biotechnology},
month = {dec},
number = {6},
pages = {1044--1053},
title = {{Microbial production of fatty acid-derived fuels and chemicals}},
volume = {24},
year = {2013}
}
@article{Chi2011,
author = {Chi, Xiaoyuan and Yang, Qingli and Pan, Lijuan and Chen, Mingna and He, Yanan and Yang, Zhen and Yu, Shanlin},
doi = {10.1007/s00299-011-1048-4},
issn = {0721-7714},
journal = {Plant Cell Reports},
month = {aug},
number = {8},
pages = {1393--1404},
title = {{Isolation and characterization of fatty acid desaturase genes from peanut (Arachis hypogaea L.)}},
volume = {30},
year = {2011}
}
@article{Dong2016,
author = {Dong, Chun-Juan and Cao, Ning and Zhang, Zhi-Gang and Shang, Qing-Mao},
doi = {10.1371/journal.pone.0149917},
editor = {Shi, Haitao},
issn = {1932-6203},
journal = {PLOS ONE},
month = {mar},
number = {3},
pages = {e0149917},
title = {{Characterization of the Fatty Acid Desaturase Genes in Cucumber: Structure, Phylogeny, and Expression Patterns}},
volume = {11},
year = {2016}
}
@article{Garba2018,
abstract = {Membrane-bound fatty acid desaturases perform oxygenated desaturation reactions to insert double bonds within fatty acyl chains in regioselective and stereoselective manners. The $\Delta$9-fatty acid desaturase strictly creates the first double bond between C9 and 10 positions of most saturated substrates. As the three-dimensional structures of the bacterial membrane fatty acid desaturases are not available, relevant information about the enzymes are derived from their amino acid sequences, site-directed mutagenesis and domain swapping in similar membrane-bound desaturases. The cold-tolerant Pseudomonas sp. AMS8 was found to produce high amount of monounsaturated fatty acids at low temperature. Subsequently, an active $\Delta$9-fatty acid desaturase was isolated and functionally expressed in Escherichia coli . In this paper we report homology modeling and docking studies of a $\Delta$9-fatty acid desaturase from a Cold-tolerant Pseudomonas sp. AMS8 for the first time to the best of our knowledge. Three dimensional structure of the enzyme was built using MODELLER version 9.18 using a suitable template. The protein model contained the three conserved-histidine residues typical for all membrane-bound desaturase catalytic activity. The structure was subjected to energy minimization and checked for correctness using Ramachandran plots and ERRAT, which showed a good quality model of 91.6 and 65.0\%, respectively. The protein model was used to preform MD simulation and docking of palmitic acid using CHARMM36 force field in GROMACS Version 5 and Autodock tool Version 4.2, respectively. The docking simulation with the lowest binding energy, −6.8 kcal/mol had a number of residues in close contact with the docked palmitic acid namely, Ile26, Tyr95, Val179, Gly180, Pro64, Glu203, His34, His206, His71, Arg182, Thr85, Lys98 and His177. Interestingly, among the binding residues are His34, His71 and His206 from the first, second, and third conserved histidine motif, respectively, which constitute the active site of the enzyme. The results obtained are in compliance with the in vivo activity of the $\Delta$9-fatty acid desaturase on the membrane phospholipids.},
author = {Garba, Lawal and {Mohamad Yussoff}, Mohamad Ariff and {Abd Halim}, Khairul Bariyyah and Ishak, Siti Nor Hasmah and {Mohamad Ali}, Mohd Shukuri and Oslan, Siti Nurbaya and {Raja Abd. Rahman}, Raja Noor Zaliha},
doi = {10.7717/peerj.4347},
issn = {2167-8359},
journal = {PeerJ},
month = {mar},
pages = {e4347},
title = {{Homology modeling and docking studies of a {$\Delta$}9-fatty acid desaturase from a Cold-tolerant Pseudomonas sp. AMS8}},
volume = {6},
year = {2018}
}
@article{Klebe2013,
author = {Klebe, Gerhard},
doi = {10.1007/978-3-642-17907-5\_13},
journal = {Drug Design},
pages = {265-290},
publisher = {Springer, Berlin, Heidelberg},
title = {Experimental Methods of Structure Determination},
year = {2013}
}
@book{bancroft1973mossbauer,
  title={M{\"o}ssbauer spectroscopy: an introduction for inorganic chemists and geochemists},
  author={Bancroft, George Michael},
  year={1973},
  publisher={John Wiley \& Sons}
}
@incollection{Sadler1989,
author = {Sadler, David M.},
booktitle = {Comprehensive Polymer Science and Supplements},
doi = {10.1016/B978-0-08-096701-1.00032-X},
pages = {731--763},
publisher = {Elsevier},
title = {{Neutron Scattering from Solid Polymers}},
year = {1989}
}
@article{Castellanos2017,
author = {Castellanos, Maria Monica and McAuley, Arnold and Curtis, Joseph E.},
doi = {10.1016/j.csbj.2016.12.004},
issn = {20010370},
journal = {Computational and Structural Biotechnology Journal},
pages = {117--130},
title = {{Investigating Structure and Dynamics of Proteins in Amorphous Phases Using Neutron Scattering}},
volume = {15},
year = {2017}
}
@article{Lindqvist1996,
abstract = {The three-dimensional structure of recombinant homodimeric delta9 stearoyl-acyl carrier protein desaturase, the archetype of the soluble plant fatty acid desaturases that convert saturated to unsaturated fatty acids, has been determined by protein crystallographic methods to a resolution of 2.4 angstroms. The structure was solved by a combination of single isomorphous replacement, anomalous contribution from the iron atoms to the native diffraction data and 6-fold non-crystallographic symmetry averaging. The 363 amino acid monomer consists of a single domain of 11 alpha-helices. Nine of these form an antiparallel helix bundle. The enzyme subunit contains a di-iron centre, with ligands from four of the alpha-helices in the helix bundle. The iron ions are bound in a highly symmetric environment, with one of the irons forming interactions with the side chains of E196 and H232 and the second iron with the side chains of E105 and H146. Two additional glutamic acid side chains, from E143 and E229, are within coordination distance to both iron ions. A water molecule is found within the second coordination sphere from the iron atoms. The lack of electron density corresponding to a mu-oxo bridge, and the long (4.2 angstroms) distance between the iron ions suggests that this probably represents the diferrous form of the enzyme. A deep channel which probably binds the fatty acid extends from the surface into the interior of the enzyme. Modelling of the substrate, stearic acid, into this channel places the delta9 carbon atom in the vicinity of one of the iron ions.},
archivePrefix = {arXiv},
arxivId = {10.2210/pdb1afr/pdb},
author = {Lindqvist, Y and Huang, W and Schneider, G and Shanklin, J},
doi = {8861937},
issn = {0261-4189},
journal = {The EMBO journal},
month = {aug},
number = {16},
pages = {4081--92},
pmid = {8861937},
primaryClass = {10.2210},
title = {{Crystal structure of delta9 stearoyl-acyl carrier protein desaturase from castor seed and its relationship to other di-iron proteins.}},
volume = {15},
year = {1996}
}
@article{Dyer2005,
abstract = {Genome sequencing showed that two proteins in Mycobacterium tuberculosis H37Rv contain the metal binding motif (D/E)X(2)HX(approximately 100)(D/E)X(2)H characteristic of the soluble diiron enzyme superfamily. These putative acyl-ACP desaturase genes desA1 and desA2 were cloned from genomic DNA and expressed in Escherichia coli BL21(DE3). DesA1 was found to be insoluble, but in contrast, DesA2 was a soluble protein amenable to biophysical characterization. Here, we report the 2.0 A resolution X-ray structure of DesA2 determined by multiple anomalous dispersion (MAD) phasing from a Se-met derivative and refinement against diffraction data obtained on the native protein. The X-ray structure shows that DesA2 is a homodimeric protein with a four-helix bundle core flanked by five additional helices that overlay with 192 structurally equivalent amino acids in the structure of stearoyl-ACP Delta9 desaturase from castor plant with an rms difference 1.42 A. In the DesA2 crystals, one metal (likely Mn from the crystallization buffer) was bound in high occupancy at the B-site of the conserved metal binding motif, while the A-site was not occupied by a metal ion. Instead, the amino group of Lys-76 occupied this position. The relationships between DesA2 and known diiron enzymes are discussed.},
author = {Dyer, David H and Lyle, Karen S and Rayment, Ivan and Fox, Brian G},
doi = {10.1110/ps.041288005},
issn = {0961-8368},
journal = {Protein science : a publication of the Protein Society},
month = {jun},
number = {6},
pages = {1508--17},
pmid = {15929999},
title = {{X-ray structure of putative acyl-ACP desaturase DesA2 from Mycobacterium tuberculosis H37Rv.}},
volume = {14},
year = {2005}
}
@article{Dar2017,
author = {Dar, Aejaz A. and Choudhury, Abhikshit R. and Kancharla, Pavan K. and Arumugam, Neelakantan},
doi = {10.3389/fpls.2017.01789},
issn = {1664-462X},
journal = {Frontiers in Plant Science},
month = {oct},
title = {{The FAD2 Gene in Plants: Occurrence, Regulation, and Role}},
volume = {8},
year = {2017}
}
@article{Madeira2019,
abstract = {The EMBL-EBI provides free access to popular bioinformatics sequence analysis applications as well as to a full-featured text search engine with powerful cross-referencing and data retrieval capabilities. Access to these services is provided via user-friendly web interfaces and via established RESTful and SOAP Web Services APIs (https://www.ebi.ac.uk/seqdb/confluence/display/JDSAT/EMBL-EBI+Web+Services+APIs+-+Data+Retrieval). Both systems have been developed with the same core principles that allow them to integrate an ever-increasing volume of biological data, making them an integral part of many popular data resources provided at the EMBL-EBI. Here, we describe the latest improvements made to the frameworks which enhance the interconnectivity between public EMBL-EBI resources and ultimately enhance biological data discoverability, accessibility, interoperability and reusability.},
author = {Madeira, F{\'{a}}bio and mi Park, Young and Lee, Joon and Buso, Nicola and Gur, Tamer and Madhusoodanan, Nandana and Basutkar, Prasad and Tivey, Adrian R N and Potter, Simon C and Finn, Robert D and Lopez, Rodrigo},
doi = {10.1093/nar/gkz268},
issn = {0305-1048},
journal = {Nucleic Acids Research},
month = {jul},
number = {W1},
pages = {W636--W641},
title = {{The EMBL-EBI search and sequence analysis tools APIs in 2019}},
volume = {47},
year = {2019}
}
@article{Apra2020,
author = {Apr{\`{a}}, E. and Bylaska, E. J. and de Jong, W. A. and Govind, N. and Kowalski, K. and Straatsma, T. P. and Valiev, M. and van Dam, H. J. J. and Alexeev, Y. and Anchell, J. and Anisimov, V. and Aquino, F. W. and Atta-Fynn, R. and Autschbach, J. and Bauman, N. P. and Becca, J. C. and Bernholdt, D. E. and Bhaskaran-Nair, K. and Bogatko, S. and Borowski, P. and Boschen, J. and Brabec, J. and Bruner, A. and Cau{\"{e}}t, E. and Chen, Y. and Chuev, G. N. and Cramer, C. J. and Daily, J. and Deegan, M. J. O. and Dunning, T. H. and Dupuis, M. and Dyall, K. G. and Fann, G. I. and Fischer, S. A. and Fonari, A. and Fr{\"{u}}chtl, H. and Gagliardi, L. and Garza, J. and Gawande, N. and Ghosh, S. and Glaesemann, K. and G{\"{o}}tz, A. W. and Hammond, J. and Helms, V. and Hermes, E. D. and Hirao, K. and Hirata, S. and Jacquelin, M. and Jensen, L. and Johnson, B. G. and J{\'{o}}nsson, H. and Kendall, R. A. and Klemm, M. and Kobayashi, R. and Konkov, V. and Krishnamoorthy, S. and Krishnan, M. and Lin, Z. and Lins, R. D. and Littlefield, R. J. and Logsdail, A. J. and Lopata, K. and Ma, W. and Marenich, A. V. and {Martin del Campo}, J. and Mejia-Rodriguez, D. and Moore, J. E. and Mullin, J. M. and Nakajima, T. and Nascimento, D. R. and Nichols, J. A. and Nichols, P. J. and Nieplocha, J. and Otero-de-la-Roza, A. and Palmer, B. and Panyala, A. and Pirojsirikul, T. and Peng, B. and Peverati, R. and Pittner, J. and Pollack, L. and Richard, R. M. and Sadayappan, P. and Schatz, G. C. and Shelton, W. A. and Silverstein, D. W. and Smith, D. M. A. and Soares, T. A. and Song, D. and Swart, M. and Taylor, H. L. and Thomas, G. S. and Tipparaju, V. and Truhlar, D. G. and Tsemekhman, K. and {Van Voorhis}, T. and V{\'{a}}zquez-Mayagoitia, {\'{A}}. and Verma, P. and Villa, O. and Vishnu, A. and Vogiatzis, K. D. and Wang, D. and Weare, J. H. and Williamson, M. J. and Windus, T. L. and Woli{\'{n}}ski, K. and Wong, A. T. and Wu, Q. and Yang, C. and Yu, Q. and Zacharias, M. and Zhang, Z. and Zhao, Y. and Harrison, R. J.},
doi = {10.1063/5.0004997},
issn = {0021-9606},
journal = {The Journal of Chemical Physics},
month = {may},
number = {18},
pages = {184102},
title = {{NWChem: Past, present, and future}},
volume = {152},
year = {2020}
}
@article{Lindahl2021,
author = {Lindahl and Abraham and Hess and van der Spoel},
doi = {10.5281/ZENODO.4723561},
keywords = {Software MD Documentation},
month = {may},
title = {{GROMACS 2021.2 Manual}},
year = {2021}
}
@article{Hess2008,
author = {Hess, Berk},
doi = {10.1021/ct700200b},
issn = {1549-9618},
journal = {Journal of Chemical Theory and Computation},
month = {jan},
number = {1},
pages = {116--122},
title = {{P-LINCS: A Parallel Linear Constraint Solver for Molecular Simulation}},
volume = {4},
year = {2008}
}
@article{Miyamoto1992,
author = {Miyamoto, Shuichi and Kollman, Peter A.},
doi = {10.1002/jcc.540130805},
issn = {0192-8651},
journal = {Journal of Computational Chemistry},
month = {oct},
number = {8},
pages = {952--962},
title = {{Settle: An analytical version of the SHAKE and RATTLE algorithm for rigid water models}},
volume = {13},
year = {1992}
}
@article{Bussi2007,
author = {Bussi, Giovanni and Donadio, Davide and Parrinello, Michele},
doi = {10.1063/1.2408420},
issn = {0021-9606},
journal = {The Journal of Chemical Physics},
month = {jan},
number = {1},
pages = {014101},
title = {{Canonical sampling through velocity rescaling}},
volume = {126},
year = {2007}
}
@article{Lindorff-Larsen2010,
author = {Lindorff-Larsen, Kresten and Piana, Stefano and Palmo, Kim and Maragakis, Paul and Klepeis, John L. and Dror, Ron O. and Shaw, David E.},
doi = {10.1002/prot.22711},
issn = {08873585},
journal = {Proteins: Structure, Function, and Bioinformatics},
month = {jun},
number = {8},
pages = {1950--1958},
title = {{Improved side-chain torsion potentials for the Amber ff99SB protein force field}},
volume = {78},
year = {2010}
}
@article{Wang2004,
author = {Wang, Junmei and Wolf, Romain M. and Caldwell, James W. and Kollman, Peter A. and Case, David A.},
doi = {10.1002/jcc.20035},
issn = {0192-8651},
journal = {Journal of Computational Chemistry},
month = {jul},
number = {9},
pages = {1157--1174},
title = {{Development and testing of a general amber force field}},
volume = {25},
year = {2004}
}
@article{Essmann1995,
author = {Essmann, Ulrich and Perera, Lalith and Berkowitz, Max L. and Darden, Tom and Lee, Hsing and Pedersen, Lee G.},
doi = {10.1063/1.470117},
issn = {0021-9606},
journal = {The Journal of Chemical Physics},
month = {nov},
number = {19},
pages = {8577--8593},
title = {{A smooth particle mesh Ewald method}},
volume = {103},
year = {1995}
}
@article{Jorgensen1983,
author = {Jorgensen, William L. and Chandrasekhar, Jayaraman and Madura, Jeffry D. and Impey, Roger W. and Klein, Michael L.},
doi = {10.1063/1.445869},
issn = {0021-9606},
journal = {The Journal of Chemical Physics},
month = {jul},
number = {2},
pages = {926--935},
title = {{Comparison of simple potential functions for simulating liquid water}},
volume = {79},
year = {1983}
}
@article{Joung2008,
author = {Joung, In Suk and Cheatham, Thomas E.},
doi = {10.1021/jp8001614},
issn = {1520-6106},
journal = {The Journal of Physical Chemistry B},
month = {jul},
number = {30},
pages = {9020--9041},
title = {{Determination of Alkali and Halide Monovalent Ion Parameters for Use in Explicitly Solvated Biomolecular Simulations}},
volume = {112},
year = {2008}
}
@unpublished{PyMOL,
	Annote = {PyMOL

The PyMOL Molecular Graphics System, Version 1.8, Schr{\"o}dinger, LLC.},
	Author = {{Schr\"odinger, LLC}},
	Date-Added = {2010-08-19 17:29:55 -0400},
	Date-Modified = {2015-12-22 18:04:08 -0400},
	Month = {November},
	Title = {The {PyMOL} Molecular Graphics System, Version~1.8},
	Year = {2015}}
@article{Maier2015,
author = {Maier, James A. and Martinez, Carmenza and Kasavajhala, Koushik and Wickstrom, Lauren and Hauser, Kevin E. and Simmerling, Carlos},
doi = {10.1021/acs.jctc.5b00255},
issn = {1549-9618},
journal = {Journal of Chemical Theory and Computation},
month = {aug},
number = {8},
pages = {3696--3713},
title = {{ff14SB: Improving the Accuracy of Protein Side Chain and Backbone Parameters from ff99SB}},
volume = {11},
year = {2015}
}
@article{Berendsen1984,
author = {Berendsen, H. J. C. and Postma, J. P. M. and van Gunsteren, W. F. and DiNola, A. and Haak, J. R.},
doi = {10.1063/1.448118},
issn = {0021-9606},
journal = {The Journal of Chemical Physics},
month = {oct},
number = {8},
pages = {3684--3690},
title = {{Molecular dynamics with coupling to an external bath}},
volume = {81},
year = {1984}
}
@article{Parrinello1981,
author = {Parrinello, M. and Rahman, A.},
doi = {10.1063/1.328693},
issn = {0021-8979},
journal = {Journal of Applied Physics},
month = {dec},
number = {12},
pages = {7182--7190},
title = {{Polymorphic transitions in single crystals: A new molecular dynamics method}},
volume = {52},
year = {1981}
}
@article{Nose1983,
author = {Nos{\'{e}}, Shuichi and Klein, M.L.},
doi = {10.1080/00268978300102851},
issn = {0026-8976},
journal = {Molecular Physics},
month = {dec},
number = {5},
pages = {1055--1076},
title = {{Constant pressure molecular dynamics for molecular systems}},
volume = {50},
year = {1983}
}
@article{Bayly1993,
author = {Bayly, Christopher I. and Cieplak, Piotr and Cornell, Wendy and Kollman, Peter A.},
doi = {10.1021/j100142a004},
issn = {0022-3654},
journal = {The Journal of Physical Chemistry},
month = {oct},
number = {40},
pages = {10269--10280},
title = {{A well-behaved electrostatic potential based method using charge restraints for deriving atomic charges: the RESP model}},
volume = {97},
year = {1993}
}
@article{Jumper2021,
abstract = {Proteins are essential to life, and understanding their structure can facilitate a mechanistic understanding of their function. Through an enormous experimental effort 1–4 , the structures of around 100,000 unique proteins have been determined 5 , but this represents a small fraction of the billions of known protein sequences 6,7 . Structural coverage is bottlenecked by the months to years of painstaking effort required to determine a single protein structure. Accurate computational approaches are needed to address this gap and to enable large-scale structural bioinformatics. Predicting the three-dimensional structure that a protein will adopt based solely on its amino acid sequence—the structure prediction component of the ‘protein folding problem' 8 —has been an important open research problem for more than 50 years 9 . Despite recent progress 10–14 , existing methods fall far short of atomic accuracy, especially when no homologous structure is available. Here we provide the first computational method that can regularly predict protein structures with atomic accuracy even in cases in which no similar structure is known. We validated an entirely redesigned version of our neural network-based model, AlphaFold, in the challenging 14th Critical Assessment of protein Structure Prediction (CASP14) 15 , demonstrating accuracy competitive with experimental structures in a majority of cases and greatly outperforming other methods. Underpinning the latest version of AlphaFold is a novel machine learning approach that incorporates physical and biological knowledge about protein structure, leveraging multi-sequence alignments, into the design of the deep learning algorithm.},
author = {Jumper, John and Evans, Richard and Pritzel, Alexander and Green, Tim and Figurnov, Michael and Ronneberger, Olaf and Tunyasuvunakool, Kathryn and Bates, Russ and {\v{Z}}{\'{i}}dek, Augustin and Potapenko, Anna and Bridgland, Alex and Meyer, Clemens and Kohl, Simon A. A. and Ballard, Andrew J. and Cowie, Andrew and Romera-Paredes, Bernardino and Nikolov, Stanislav and Jain, Rishub and Adler, Jonas and Back, Trevor and Petersen, Stig and Reiman, David and Clancy, Ellen and Zielinski, Michal and Steinegger, Martin and Pacholska, Michalina and Berghammer, Tamas and Bodenstein, Sebastian and Silver, David and Vinyals, Oriol and Senior, Andrew W. and Kavukcuoglu, Koray and Kohli, Pushmeet and Hassabis, Demis},
doi = {10.1038/s41586-021-03819-2},
issn = {0028-0836},
journal = {Nature},
month = {jul},
title = {{Highly accurate protein structure prediction with AlphaFold}},
year = {2021}
}
@article{Ye2003,
author = {Ye, Y. and Godzik, A.},
doi = {10.1093/bioinformatics/btg1086},
issn = {1367-4803},
journal = {Bioinformatics},
month = {sep},
number = {Suppl 2},
pages = {ii246--ii255},
title = {{Flexible structure alignment by chaining aligned fragment pairs allowing twists}},
volume = {19},
year = {2003}
}
@article{Li2020,
abstract = {FATCAT 2.0 server (http://fatcat.godziklab.org/), provides access to a flexible protein structure alignment algorithm developed in our group. In such an alignment, rotations and translations between elements in the structure are allowed to minimize the overall root mean square deviation (RMSD) between the compared structures. This allows to effectively compare protein structures even if they underwent structural rearrangements in different functional forms, different crystallization conditions or as a result of mutations. The major update for the server introduces a new graphical interface, much faster database searches and several new options for visualization of the structural differences between proteins},
author = {Li, Zhanwen and Jaroszewski, Lukasz and Iyer, Mallika and Sedova, Mayya and Godzik, Adam},
doi = {10.1093/nar/gkaa443},
issn = {0305-1048},
journal = {Nucleic Acids Research},
month = {jul},
number = {W1},
pages = {W60--W64},
title = {{FATCAT 2.0: towards a better understanding of the structural diversity of proteins}},
volume = {48},
year = {2020}
}
@misc{Uttaro2006,
abstract = {Polyunsaturated fatty acids have important structural roles in cell membranes. They are also intermediates in the synthesis of biologically active molecules such as eicosanoids, which mediate fever, inflammation, blood pressure and neurotransmission. Arachidonic and docosahexaenoic acids are essential components of brain tissues and, through their involvement in the development of neural and retinal functions, important dietary nutrients for neonatal babies. Lower eukaryotes are particularly rich in C20-22 polyunsaturated fatty acids. Fungi and marine microalgae are currently used to produce nutraceutic oils. Other protists and algae are being studied because of the variability in their enzymes involved in polyunsaturated fatty acid biosynthesis. Such enzymes could be used as source for the production of transgenic organisms able to synthesize designed oils for human diet or, in the case of parasitic protozoa, they might be identified as putative chemotherapeutic targets. Polyunsaturated fatty acids can be synthesized by two different pathways: an anaerobic one, by using polyketide synthase related enzymes, and an aerobic one, which involves the action of elongases and oxygen dependent desaturases. Desaturases can be classified into three main types, depending on which of the consecutive steps of polyunsaturated fatty acid synthesis they are involved with. The enzymes may be specialized to act on: saturated substrates (type I); mono- and di-unsaturated fatty acids by introducing additional double bonds at the methyl-end site of the existing double bonds (type II); or the carboxy half ('front-end') of polyunsaturated ones (type III). Type III desaturases require the alternating action of elongases. A description of the enzymes that have been isolated and functionally characterized is provided, in order to highlight the different pathways found in lower eukaryotes. {\textcopyright} 2006 IUBMB.},
author = {Uttaro, Antonio D.},
booktitle = {IUBMB Life},
doi = {10.1080/15216540600920899},
issn = {15216543},
keywords = {Algae,Desaturases,Elongases,Fungi,Polyunsaturated fatty acids,Protozoa,Trypanosomatids},
month = {oct},
number = {10},
pages = {563--571},
pmid = {17028087},
title = {{Biosynthesis of polyunsaturated fatty acids in lower eukaryotes}},
volume = {58},
year = {2006}
}
@article{Shindyalov1998,
author = {Shindyalov, I. N. and Bourne, P. E.},
doi = {10.1093/protein/11.9.739},
issn = {1741-0126},
journal = {Protein Engineering Design and Selection},
month = {sep},
number = {9},
pages = {739--747},
title = {{Protein structure alignment by incremental combinatorial extension (CE) of the optimal path}},
volume = {11},
year = {1998}
}
@article{Holm2020,
author = {Holm, Liisa},
doi = {10.1002/pro.3749},
issn = {0961-8368},
journal = {Protein Science},
month = {jan},
number = {1},
pages = {128--140},
title = {{DALI and the persistence of protein shape}},
volume = {29},
year = {2020}
}
@article{Hertig2016,
author = {Hertig, Samuel and Latorraca, Naomi R. and Dror, Ron O.},
doi = {10.1371/journal.pcbi.1004746},
editor = {Liu, Jin},
issn = {1553-7358},
journal = {PLOS Computational Biology},
month = {jun},
number = {6},
pages = {e1004746},
title = {{Revealing Atomic-Level Mechanisms of Protein Allostery with Molecular Dynamics Simulations}},
volume = {12},
year = {2016}
}
@article{Ferro2003,
author = {Ferro, Myriam and Salvi, Daniel and Brugi{\`{e}}re, Sabine and Miras, St{\'{e}}phane and Kowalski, Sol{\`{e}}ne and Louwagie, Mathilde and Garin, J{\'{e}}r{\^{o}}me and Joyard, Jacques and Rolland, Norbert},
doi = {10.1074/mcp.M300030-MCP200},
issn = {15359476},
journal = {Molecular {\&} Cellular Proteomics},
month = {may},
number = {5},
pages = {325--345},
title = {{Proteomics of the Chloroplast Envelope Membranes from Arabidopsis thaliana}},
url = {https://linkinghub.elsevier.com/retrieve/pii/S1535947620348611},
volume = {2},
year = {2003}
}
@article{Chen2013,
author = {Chen, Haiqin and Gu, Zhennan and Zhang, Hao and Wang, Mingxuan and Chen, Wei and Lowther, W. Todd and Chen, Yong Q.},
doi = {10.1371/journal.pone.0058139},
editor = {Schunck, Wolf-Hagen},
issn = {1932-6203},
journal = {PLoS ONE},
month = {mar},
number = {3},
pages = {e58139},
title = {{Expression and Purification of Integral Membrane Fatty Acid Desaturases}},
url = {https://dx.plos.org/10.1371/journal.pone.0058139},
volume = {8},
year = {2013}
}
@article{Baek2021,
abstract = {DeepMind presented remarkably accurate predictions at the recent CASP14 protein structure prediction assessment conference. We explored network architectures incorporating related ideas and obtained the best performance with a three-track network in which information at the 1D sequence level, the 2D distance map level, and the 3D coordinate level is successively transformed and integrated. The three-track network produces structure predictions with accuracies approaching those of DeepMind in CASP14, enables the rapid solution of challenging X-ray crystallography and cryo-EM structure modeling problems, and provides insights into the functions of proteins of currently unknown structure. The network also enables rapid generation of accurate protein-protein complex models from sequence information alone, short circuiting traditional approaches which require modeling of individual subunits followed by docking. We make the method available to the scientific community to speed biological research.},
author = {Baek, Minkyung and DiMaio, Frank and Anishchenko, Ivan and Dauparas, Justas and Ovchinnikov, Sergey and Lee, Gyu Rie and Wang, Jue and Cong, Qian and Kinch, Lisa N. and Schaeffer, R. Dustin and Mill{\'{a}}n, Claudia and Park, Hahnbeom and Adams, Carson and Glassman, Caleb R. and DeGiovanni, Andy and Pereira, Jose H. and Rodrigues, Andria V. and van Dijk, Alberdina A. and Ebrecht, Ana C. and Opperman, Diederik J. and Sagmeister, Theo and Buhlheller, Christoph and Pavkov-Keller, Tea and Rathinaswamy, Manoj K. and Dalwadi, Udit and Yip, Calvin K. and Burke, John E. and Garcia, K. Christopher and Grishin, Nick V. and Adams, Paul D. and Read, Randy J. and Baker, David},
doi = {10.1126/science.abj8754},
issn = {0036-8075},
journal = {Science},
month = {jul},
pages = {eabj8754},
title = {{Accurate prediction of protein structures and interactions using a three-track neural network}},
url = {https://www.sciencemag.org/lookup/doi/10.1126/science.abj8754},
year = {2021}
}
@article{Paysan-Lafosse2023,
abstract = {The InterPro database (https://www.ebi.ac.uk/interpro/) provides an integrative classification of protein sequences into families, and identifies functionally important domains and conserved sites. Here, we report recent developments with InterPro (version 90.0) and its associated software, including updates to data content and to the website. These developments extend and enrich the information provided by InterPro, and provide a more user friendly access to the data. Additionally, we have worked on adding Pfam website features to the InterPro website, as the Pfam website will be retired in late 2022. We also show that InterPro's sequence coverage has kept pace with the growth of UniProtKB. Moreover, we report the development of a card game as a method of engaging the non-scientific community. Finally, we discuss the benefits and challenges brought by the use of artificial intelligence for protein structure prediction.},
author = {Paysan-Lafosse, Typhaine and Blum, Matthias and Chuguransky, Sara and Grego, Tiago and Pinto, Beatriz L{\'{a}}zaro and Salazar, Gustavo A and Bileschi, Maxwell L and Bork, Peer and Bridge, Alan and Colwell, Lucy and Gough, Julian and Haft, Daniel H and Letuni{\'{c}}, Ivica and Marchler-Bauer, Aron and Mi, Huaiyu and Natale, Darren A and Orengo, Christine A and Pandurangan, Arun P and Rivoire, Catherine and Sigrist, Christian J A and Sillitoe, Ian and Thanki, Narmada and Thomas, Paul D and Tosatto, Silvio C E and Wu, Cathy H and Bateman, Alex},
doi = {10.1093/nar/gkac993},
issn = {0305-1048},
journal = {Nucleic Acids Research},
month = {jan},
number = {D1},
pages = {D418--D427},
title = {{InterPro in 2022}},
url = {https://academic.oup.com/nar/article/51/D1/D418/6814474},
volume = {51},
year = {2023}
}
@article{Shanklin1994,
abstract = {The eukaryotic fatty acid desaturases are iron-containing enzymes that catalyze the NAD-(P)H- and O2-dependent introduction of double bonds into methylene-interrupted fatty acyl chains. Examination of deduced amino acid sequences for the membrane desaturases from mammals, fungi, insects, higher plants, and cyanobacteria has revealed three regions of conserved primary sequence containing HX(3 or 4)H, HX(2 or 3)HH, and HX(2 or 3)HH. This motif is also present in the bacterial membrane enzymes alkane hydroxylase ($\omega$-hydroxylase) and xylene monooxygenase. Hydropathy analyses indicate that these enzymes contain up to three long hydrophobic domains which would be long enough to span the membrane bilayer twice. The conserved His-containing regions have a consistent positioning with respect to these potential membrane spanning domains. Taken together, these observations suggest that the membrane fatty acid desaturases and hydrocarbon hydroxylases have a related protein fold, possibly arising from a common ancestral origin. In order to examine the functional role of these conserved His residues, we have made use of the ability of the rat $\Delta$9 desaturase gene to complement a yeast strain deficient in the $\Delta$9 desaturase gene function (ole1). By site-directed mutagenesis, eight conserved His residues in the rat $\Delta$9 desaturase were individually converted to Ala. Each His → Ala mutation failed to complement the yeast ole1 mutant. In contrast, mutation of three nonconserved flanking His residues or a partially conserved Arg residue within the conserved motif to Ala allowed for complementation of the ole1 phenotype. Western blot analysis showed that steady-state expression levels were equivalent for the wild-type rat desaturase and for all mutants, suggesting that the conserved His residues are essential for catalytic function. One role for these His residues would be to act as ligands for the iron atom(s) contained in these enzymes. On the basis of these results and the presently available spectroscopic data, we recognize the possibility that the membrane desaturases, alkane hydroxylase, and xylene monooxygenase contain a new structural type of diiron center. {\textcopyright} 1994, American Chemical Society. All rights reserved.},
author = {Shanklin, John and Whittle, Edward and Fox, Brian G.},
doi = {10.1021/bi00209a009},
file = {:Users/mcna892/Library/Application Support/Mendeley Desktop/Downloaded/Shanklin, Whittle, Fox - 1994 - Eight Histidine Residues Are Catalytically Essential in a Membrane-Associated Iron Enzyme, Stearoyl-CoA.pdf:pdf},
issn = {15204995},
journal = {Biochemistry},
month = {nov},
number = {43},
pages = {12787--12794},
pmid = {7947684},
title = {{Eight Histidine Residues Are Catalytically Essential in a Membrane-Associated Iron Enzyme, Stearoyl-CoA Desaturase, and Are Conserved in Alkane Hydroxylase and Xylene Monooxygenase}},
volume = {33},
year = {1994}
}
@article{Shanklin1997,
abstract = {The gene encoding the alkane $\omega$-hydroxylase (AlkB; EC 1.14.15.3 ) from Pseudomonas oleovorans was expressed in Escherichia coli . The integral-membrane protein was purified as nearly homogeneous protein vesicles by differential ultracentrifugation and HPLC cation exchange chromatography without the detergent solubilization normally required for membrane proteins. Purified AlkB had specific activity of up to 5 units/mg for octane-dependent NADPH consumption. M{\"{o}}ssbauer studies of AlkB showed that it contains an exchange-coupled dinuclear iron cluster of the type found in soluble diiron proteins such as hemerythrin, ribonucleotide reductase, methane monooxygenase, stearoyl-acyl carrier protein (ACP) $\Delta$ 9 desaturase, rubrerythrin, and purple acid phosphatase. In the as-isolated enzyme, the cluster contains an antiferromagnetically coupled pair of high-spin Fe(III) sites, with an occupancy of up to 0.9 cluster per AlkB. The diferric cluster could be reduced by sodium dithionite, and the diferrous state was found to be stable in air. When both O 2 and substrate (octane) were added, however, the diferrous cluster was quantitatively reoxidized, proving that the diiron cluster occupies the active site. M{\"{o}}ssbauer data on reduced AlkB are consistent with a cluster coordination rich in nitrogen-containing ligands. New sequence analyses indicate that at least 11 nonheme integral-membrane enzymes, including AlkB, contain the 8-histidine motif required for catalytic activity in stearoyl-CoA desaturase. Based on our M{\"{o}}ssbauer studies of AlkB, we propose that the integral-membrane enzymes in this family contain diiron clusters. Because these enzymes catalyze a diverse range of oxygenation reactions, this proposal suggests a greatly expanded role for diiron clusters in O 2 -activation biochemistry.},
author = {Shanklin, John and Achim, Catalina and Schmidt, Hermann and Fox, Brian G. and M{\"{u}}nck, Eckard},
doi = {10.1073/pnas.94.7.2981},
issn = {0027-8424},
journal = {Proceedings of the National Academy of Sciences},
month = {apr},
number = {7},
pages = {2981--2986},
title = {{M{\"{o}}ssbauer studies of alkane $\omega$-hydroxylase: Evidence for a diiron cluster in an integral-membrane enzyme}},
url = {https://pnas.org/doi/full/10.1073/pnas.94.7.2981},
volume = {94},
year = {1997}
}
@article{Cai2020,
author = {Cai, Yuanheng and Yu, Xiao-Hong and Chai, Jin and Liu, Chang-Jun and Shanklin, John},
doi = {10.1074/jbc.RA120.014258},
issn = {00219258},
journal = {Journal of Biological Chemistry},
month = {aug},
number = {32},
pages = {11337--11345},
title = {{A conserved evolutionary mechanism permits $\Delta$9 desaturation of very-long-chain fatty acyl lipids}},
url = {https://linkinghub.elsevier.com/retrieve/pii/S0021925817492234},
volume = {295},
year = {2020}
}
@article{Cai2018,
author = {Cai, Yuanheng and Yu, Xiao-Hong and Liu, Qun and Liu, Chang-Jun and Shanklin, John},
doi = {10.1074/jbc.RA118.005972},
issn = {00219258},
journal = {Journal of Biological Chemistry},
month = {dec},
number = {51},
pages = {19844--19853},
title = {{Two clusters of residues contribute to the activity and substrate specificity of Fm1, a bifunctional oleate and linoleate desaturase of fungal origin}},
url = {https://linkinghub.elsevier.com/retrieve/pii/S0021925820311108},
volume = {293},
year = {2018}
}
