## Supplementary material for "Data-Driven Modeling and Analysis of Fatty Acid Desaturase in Plants": TeX Files: paper.tex

%% 
%% Copyright 2007-2020 Elsevier Ltd
%% 
%% This file is part of the 'Elsarticle Bundle'.
%% ---------------------------------------------
%% 
%% It may be distributed under the conditions of the LaTeX Project Public
%% License, either version 1.2 of this license or (at your option) any
%% later version.  The latest version of this license is in
%%    http://www.latex-project.org/lppl.txt
%% and version 1.2 or later is part of all distributions of LaTeX
%% version 1999/12/01 or later.
%% 
%% The list of all files belonging to the 'Elsarticle Bundle' is
%% given in the file `manifest.txt'.
%% 
%% Template article for Elsevier's document class `elsarticle'
%% with harvard style bibliographic references

\documentclass[preprint,12pt]{elsarticle}

%% Use the option review to obtain double line spacing
%% \documentclass[preprint,review,12pt]{elsarticle}

%% Use the options 1p,twocolumn; 3p; 3p,twocolumn; 5p; or 5p,twocolumn
%% for a journal layout:
%% \documentclass[final,1p,times]{elsarticle}
%% \documentclass[final,1p,times,twocolumn]{elsarticle}
%% \documentclass[final,3p,times]{elsarticle}
%% \documentclass[final,3p,times,twocolumn]{elsarticle}
%% \documentclass[final,5p,times]{elsarticle}
%% \documentclass[final,5p,times,twocolumn]{elsarticle}

%% For including figures, graphicx.sty has been loaded in
%% elsarticle.cls. If you prefer to use the old commands
%% please give \usepackage{epsfig}

%% The amssymb package provides various useful mathematical symbols
\usepackage{amssymb}
%% The amsthm package provides extended theorem environments
%% \usepackage{amsthm}

%% The lineno packages adds line numbers. Start line numbering with
%% \begin{linenumbers}, end it with \end{linenumbers}. Or switch it on
%% for the whole article with \linenumbers.
%% \usepackage{lineno}

%Other Packages
%\usepackage{float}

\journal{bioRxiv}

\begin{document}

\begin{frontmatter}

%% Title, authors and addresses

%% use the tnoteref command within \title for footnotes;
%% use the tnotetext command for theassociated footnote;
%% use the fnref command within \author or \address for footnotes;
%% use the fntext command for theassociated footnote;
%% use the corref command within \author for corresponding author footnotes;
%% use the cortext command for theassociated footnote;
%% use the ead command for the email address,
%% and the form \ead[url] for the home page:
%% \title{Title\tnoteref{label1}}
%% \tnotetext[label1]{}
%% \author{Name\corref{cor1}\fnref{label2}}
%% \ead{email address}
%% \ead[url]{home page}
%% \fntext[label2]{}
%% \cortext[cor1]{}
%% \affiliation{organization={},
%%             addressline={},
%%             city={},
%%             postcode={},
%%             state={},
%%             country={}}
%% \fntext[label3]{}

\title{Data-Driven Modeling and Analysis of Fatty Acid Desaturase in Plants}
%% use optional labels to link authors explicitly to addresses:
%% \author[label1,label2]{}
%% \affiliation[label1]{organization={},
%%             addressline={},
%%             city={},
%%             postcode={},
%%             state={},
%%             country={}}
%%
%% \affiliation[label2]{organization={},
%%             addressline={},
%%             city={},
%%             postcode={},
%%             state={},
%%             country={}}

\author[inst1]{Andrew D. McNaughton}

\affiliation[inst1]{organization={Earth and Biological Science Directorate, Pacific Northwest National Laboratory},%Department and Organization
            addressline={}, 
            city={Richland},
            state={Washington},
            postcode={99352}, 
            country={United States}}
            
\author[inst2]{John Shanklin}
\affiliation[inst2]{organization={Brookhaven National Laboratory, Department of Biology},%Department and Organization
            addressline={}, 
            city={Upton},
            state={New York},
            postcode={11973}, 
            country={United States}}
            
\author[inst3]{Simone Raugei*} 

\affiliation[inst3]{organization={Physical and Computational Science Directorate, Pacific Northwest National Laboratory},%Department and Organization
            addressline={}, 
            city={Richland},
            state={Washington},
            postcode={99352}, 
            country={United States}}

\author[inst1]{Neeraj Kumar*}
\cortext[cor1]{Corresponding Authors.\newline
}
%\affiliation[inst3]

\begin{abstract}
%% Text of abstract
Fatty acids and their derivatives continue to garner attention as sustainable alternatives to petrochemically derived materials. Towards this goal, we examine the plant membrane-bound fatty acid desaturases (FADs) and develop an overview of their 3D structure and phylogenic relationships. Through this effort we developed two plant fatty acid desaturase homology models that we analyzed with substrate docking and molecular dynamics simulations to better understand the diiron binding pocket that is characteristic of these proteins. The comparison between the Omega6 and Delta12 homology models specifically in regards to the binding affinity as a function of carbon chain length indicates that there is a dip in binding affinity near the 18:2 carbon chain length but the rest maintain a rather consistent binding affinity throughout. Furthermore, MD analysis highlights the importance of regions within the cytosolic cap domain on the binding pocket.

\end{abstract}

%%Graphical abstract
\begin{graphicalabstract}
\includegraphics[scale=0.4]{FAD_GRAPHICAL_ABSTRACT.png}
\end{graphicalabstract}

%%Research highlights
\begin{highlights}
\item Developed pipeline to predict function from protein sequence 
\item Modeled two fatty acid desaturases and predicted their structure and functions
\item Leveraged docking simulations and molecular dynamics to confirm protein function
\end{highlights}

\begin{keyword}
%% keywords here, in the form: keyword \sep keyword
Fatty Acid Desaturase \sep Homology Modelling \sep FAD2 \sep FAD6 \sep diiron core \sep Fatty Acids
%% PACS codes here, in the form: \PACS code \sep code
\PACS 0000 \sep 1111
%% MSC codes here, in the form: \MSC code \sep code
%% or \MSC[2008] code \sep code (2000 is the default)
\MSC 0000 \sep 1111
\end{keyword}

\end{frontmatter}

%% \linenumbers

%% main text
\section{Introduction}
\label{sec:introduction}

%% For citations use: 
%%       \citet{<label>} ==> Jones et al. [21]
%%       \citep{<label>} ==> [21]
%%

Fatty Acid Desaturases (FADs) are a complex family of enzymes found within both prokaryotes and eukaryotes that catalyze the regiospecific dehydrogenation of fatty acids (FAs), creating a carbon–carbon double bond with the concomitant reduction of dioxygen to water \citep{Petrini2004,STUMPF1980,HARWOOD1980}. FADs show a wide range of lipid substrate preferences that includes acyl-CoAs,  sphingolipids, phospholipids, and galactolipids \citep{Sperling2003,Meesapyodsuk2012,LI2016341}. Their primary role is to preserve the structure and function of cellular membranes by utilizing the desaturation of glycerolipids that form the lipid bilayer, which typically maintains membrane fluidity \citep{Stubbs1984,Los1998}.  FADs have been explored for use in commercially relevant and sustainable green technologies, as a replacement for petroleum-derived chemicals. In addition, FAs and their derivatives have applications in the production of detergents, soaps, lubricants, cosmetics, pharmaceuticals, and biofuels \citep{Lennen2013,liu2014,Zhou2014,Yu2014,Tee2014,Runguphan2014}. However, further development of these enzymes and a better understanding of their functional role is necessary before biologically produced oils can become primary chemical feedstocks. 

There are two observed classes of FADs present in nature: soluble and integral membrane-bound desaturases. The soluble FADs are found primarily in the plastidial stroma of higher plants and are responsible for creating double bonds in FAs that are esterified to acyl carrier protein (ACP). Soluble FADs are not overly diverse and have many experimentally determined structures which have enabled a mechanistic understanding of regioselectivity \citep{Lindqvist1996,Dyer2005}. Integral membrane FADs, on the other hand, show a much  greater variation in occurrence, protein sequence, biochemical function, and substrate binding range \citep{Nachtschatt2020,Buist2004,Sperling2001,Sperling2003}, but are lacking comprehensive structural information for more accurate characterization. They are primarily located in the endomembrane system \citep{Shanklin1998} or the inner chloroplastic membrane \citep{Ferro2003}. These  membrane-bound desaturases bind iron ions with conserved histidine residues that comprise three conserved histidine boxes:  [H(X)3–4H], [H(X)2-3HH] and [H/Q(X)2–3HH] (H, histidine; X, variable amino acid; Q , glutamine)\citep{Shanklin1998,Shanklin1994}.

Currently, we have a limited understanding of the range of activity and substrate specificity for membrane-bound FADs, which makes their systematic functional characterization challenging\citep{LI2016341}. A significant problem for computational approaches is the lack of reliable tertiary (3D) structures, which prevents, for instance, the study of protein-ligand \citep{Garba2018} or protein-protein interactions \citep{Lou2014}, and consequently, drawing conclusions about protein structure-function relationships. Databases of protein sequences available through UniProt\citep{Consortium2019} and the Protein Data Bank\citep{Berman2000} offer accurate protein sequences, but do not contain accurate tertiary structures for all proteins. These structures can be resolved experimentally with a wide range of techniques \citep{Klebe2013, Bai2015, bancroft1973mossbauer, Shanklin1997, Sadler1989, Castellanos2017}, but are relatively expensive, time consuming, and often plagued by a number of difficulties, and it is unrealistic to experimentally determine every possible protein structure. To address this problem, we can leverage computational methods and tools used to predict 3D-structure from the primary protein sequence. Once a structure is predicted it can be further analyzed by  methods such as physics-based simulations and data science tools to better understand enzyme motions while also helping to predict protein function. 

\begin{figure}[!ht]
    \centering
    \includegraphics[width=\textwidth]{Screen Shot 2021-03-18 at 3.23.26 PM.png} 
    \caption{The chemical structure of common natural lipid compounds. There are two 
    common glycerol moieties at the top where X represents the phospholipid headgroup 
    and R represents the five most commonly occurring fatty acid side chains. Adapted from \citet{Nachtschatt2020}}
    \label{fig:Moieties}
\end{figure}

In FADs, substrate specificity and regioselectivity (the position the double bond is placed) is essentially determined by the interaction of the enzyme and lipid head-group (Fig \ref{fig:Moieties}). Recent  results \citep{Bai2015,Wang2015} from human and mouse stearoyl-CoA desaturase structures revealed that the hydrophilic CoA head-group from the substrate will form an electrostatic interaction and hydrogen bond with residues in the cytoplasmic domain and transmembrane helix (TM) 1 of the desaturase \citep{LI2016341}. This then causes the acyl group to orient itself into the long hydrophobic tunnel leading to the diiron active site.

Toward reliable structure-function relationships of membrane bound FADs, we collected a list of FADs with unknown structures from the higher plant \textit{Arabidopsis thaliana}, compared their sequences and phylogeny, and selected two proteins to generate homology models, whose structure was used for substrate docking and subsequent molecular dynamics simulations. Altogether these methods allow us to gather a better understanding of the underlying fatty acid side chain binding motifs and energy states in these FADs. 

\section{Results and Discussion}
\label{sec:Results}

\subsection{Data Curation of Membrane-Bound Desaturases}
\label{sec:Curate}

In order to characterize correlations between sequence and function of the membrane-bound FADs, sequences from the PFAM/InterPro \citep{pfam,Paysan-Lafosse2023} FAD family PF00487/IPR005804 were selected using the criteria laid out in previous studies \citet{LI2016341}. We selected only reviewed seed sequence clusters with a length range between 350 aa and 550 aa. In total, we obtained 71 total Arabidopsis sequences from \textbf{PFAM/InterPro}, which reduced to 10 when accounting for the length restriction and similarity scores. This selection includes $\Delta^{12}$ desaturase (P46313)(FAD2) and $\Omega^6$  chloroplastic desaturase (P46312)(FAD6), which are the target of the present study.

We took the 10 Arabidopsis enzymes (Table~\ref{tab1}) and compared their sequence alignment to identify key regions that may affect their structure-function relationship. From these 10 enzymes, we ran a multi-sequence alignment to visualize similar regions in the sequence of the different enzymes (Shown in Fig \ref{fig:sequence}. We wanted to be able to compare the histidine motifs of these proteins to see if they share key similarities common to the plant FAD family. We discovered that between these 10 enzymes, the 3-histidine box motif is conserved throughout, giving us confidence that these sequences are characteristic FAD enzymes. We also noticed the occasional replacement of the first histidine (Figure~\ref{fig:phylo}, Box 3) with a glutamate amino acid in several of the sequences.

\begin{table}[!ht]
\scriptsize
\centering
\caption{A full list of Arabidopsis fatty acid desaturase proteins and their individual 
sequence lengths, gathered from the PFAM class PF00487. 
C represents chloroplastic-bound desaturases and ER represents 
endoplasmic reticulum-bound desautrases.}

\begin{tabular}{|l|c|}
Description                                                                            & Sequence Length \\
Delta(8)-fatty-acid desaturase 2                                                       & 449             \\
Palmitoyl-monogalactosyldiacylglycerol delta-7 desaturase C                            & 371             \\
Probable lipid desaturase ADS3.2 C                                                     & 361             \\
Delta(8)-fatty-acid desaturase 1                                                       & 449             \\
Fatty acid desaturase 8                                                                & 382             \\
sn-2 acyl-lipid omega-3 desaturase (ferredoxin) C.                                     & 446             \\
Omega-6 fatty acid desaturase C                                                        & 448             \\
Delta(12)-fatty-acid desaturase                                                        & 383             \\
Temperature-sensitive sn-2 acyl-lipid omega-3 desaturase (ferredoxin) C.               & 435             \\
Acyl-lipid omega-3 desaturase (cytochrome b5) ER.                                      & 386            
\end{tabular}
\label{tab1}
\end{table}

\begin{figure}[!ht]
    \centering
    \includegraphics[width=\textwidth]{Sequence_Box.png}
    \caption{Snapshot of multiple sequence alignment of 10 Arabidopsis enzymes observed. Sequences maintain alignment with the 3 histidine box motif. Box 3 contains 2 proteins that show the ability of glutamate (Q) to replace the first histidine (H) in the motif.}
    \label{fig:sequence}
\end{figure}

\begin{figure}[!ht]
    \centering
    \includegraphics[width=\textwidth]{phylo.png}
    \caption{Phylogenic Tree analysis of the 10 Arabidopsis enzymes looked at in this study.}
    \label{fig:phylo}
\end{figure}

Figure \ref{fig:phylo} also shows the close phylogenic relationship between these 10 \textit{Arabidopsis} enzymes. Additionally, the comparison of these phylogenic relationships with the aligned sequences above showed that the two enzymes displaying the glutamate substitution in the histidine Figure~\ref{fig:phylo}, Box 3 are much more closely related phylogenically then those with the standard histidine motif.

\subsection{Homology Model of FAD2 and FAD6}

The first step toward a better characterization of FADs is the construction of 3D models for use in molecular docking simulations. To this end, we modeled the two proteins of interest: FAD2 and FAD6. No published crystal structures for FAD2 or FAD6 currently exist, so we utilized a homology-based modeling approach using PHYRE2 \citep{Kelley2015} to create a representative structural model from an existing desaturase template. The chosen 3D structure template was the crystal structure of human integral membrane stearoyl-CoA desaturase (hSCD, PDB: 4ZYO) \citep{Wang2015}, an integral membrane-bound desaturase embedded in the endoplasmic reticulum, which shares key aspects of the FAD2 and FAD6 proteins. such as a diiron core, a transmembrane region, and a central binding pocket. 

The models, labeled as AtFAD2 and AtFAD6 to represent their host organism as well as the reaction they catalyze, are shown in Fig \ref{fig:proteins}. AtFAD6 has a homology confidence level of is 99.8\% and sequence similarity is 9\%. AtFAD2 has a homology confidence of 99.9\% and sequence similarity as 14\%.

\begin{figure}[!ht]
    \centering
    \includegraphics[scale=0.8]{Homology_Full.png}
    \caption{The resulting homology models for AtFAD6 (middle, green) and AtFAD2 (right, cyan). These hold a consistent mushroom shape with 4 helix transmembrane region to the template model of hSCD (PDB: 4ZYO) (left, magenta).}
    \label{fig:proteins}
\end{figure} 

Following the results of section \ref{sec:Curate}, we know that the AtFAD2 and AtFAD6 proteins should contain the consistent histidine region surrounding the diiron core. Modeling is not able to approximate non-protein features such as iron, but the histidine motifs should still surround the location in the binding pocket where the diiron center resides. By overlaying the diiron centers from 4ZYO, we can visualize if the histidine motifs surround the correct region (shown in Fig \ref{fig:Hist}). From this snapshot we can tell that AtFAD6 is consistent with the 8-histidine motif expected of a desaturase that utilizes the diiron core as the electron donor in the $\omega$-desaturase reaction. AtFAD2 also contains 8-histidines bound to the diiron center even though it also utilizes cyt\textit{b}5 as the electron donor for $\Delta$-desaturation.

\begin{figure}[!ht]
    \centering
    \includegraphics[width=\textwidth]{Hist_Regions.png}
    \caption{Graphic overlay of the histidine regions present in the homolgoy models. The top left model is the histidine boxes present in AtFAD6 and the top right is that of AtFAD6. The bottom is an overlay of both AtFAD2 and AtFAD6 to show the structural similarities in the histidine motif.}
    \label{fig:Hist}
\end{figure}

A key takeaway from this modeling is the similarity in 3D structure maintained throughout the 2 proteins from the template. The proteins all exhibit a similar folding pattern and the catalytic cap domain on the cytosolic side of the membrane. The overall shape is quite similar as expected from the high homology confidence. Differences within the model are due to differences in the primary/secondary structure that result in a different tertiary structures. We also observe that the trans-membrane region of each protein consists of four primary helices which is consistent with the template. There is a consistent closed conformation between the homology models, showing that that each maintains the substrate entry into the binding pocket of the hSCD1 protein. The key differences amongst these structures are particularly focused in the catalytic cap domain for each protein. These differences suggest that alternative binding modes and electron transfer mechanisms may affect how the protein carries out desaturation. AtFAD6 has a catalytic domain that is primed more for dealing with 16:1 and 18:1 fatty acid desaturation whitout requiring the binding of cyt\textit{b}5 for electron donation. AtFAD2 on the other hand requires the input of electrons from cyt\textit{b}5, so its cap domain is more open to binding these structures to its surface. AtFAD2 also primarily accommodates the desaturation of 18:1 fatty acid, so the binding pocket may be optimized for this process. Each protein has a similarly located diiron center off of the cytosolic plane, without much variation between them. One limitation with the homology modeling process, however, is that it does not seem to account for residues 158-179 of AtFAD2 and attempts to bridge residue 157 to residue 180 directly. This abridged segment of the structure could be why the caps of each modeled protein appear different.

\subsection{Substrate Docking}
\label{sec:Docking}
A key follow up to homology modeling is how the model accepts and binds substrates to the structure. In this capacity, we utilized docking of various carbon chain length fatty acid substrates from the range of C12 to C26 in both a blind docking configuration and in the vicinity of the models binding pocket. To ensure we best modeled the structure of the target FAs, we utilized DFT calculations as implemented in NWChem \cite{Apra2020} to obtain quantum mechanically optimal features for the docking. 

A key result we want to visualize is the location of the docked proteins once the allowed docking region was refined. Fig \ref{fig:Channel} below highlights the results of docking in the binding pocket around the diiron center of AtFAD6 and AtFAD2. This allows us to see that the FAs are in fact binding in the expected binding pocket within the protein surface.

\begin{figure}[!ht]
    \centering
    \includegraphics[width=\textwidth]{Binding_Channels.png}
    \caption{Homology model of AtFAD6 (left) and AtFAD2 (right) with binding pocket highlighted. These figures show where the expected desaturase substrates are binding within the central active site surrounding the diiron core. Palmitoleic acid (16:1 fatty acid) is shown as the substrate for AtFAD6 and Oleic acid (18:1 fatty acid) is shown as the substrate for AtFAD2.}
    \label{fig:Channel}
\end{figure} 

An important result of this is the effect of carbon chain length on the binding affinity. We plot this effect to visualize this relationship in Figure \ref{fig:affinity}.

\begin{figure}[!ht]
    \centering
    \includegraphics[width=\textwidth]{FAD_BA_STACK.png}
    \caption{A comparison between the Omega6 and Delta12 homology models specifically in regards to the binding affinity (kcal/mol) as a function of carbon chain length. We see that there is a dip in binding affinity near the 18:2 carbon chain length but the rest maintain a rather consistent binding affinity throughout. Chain lengths are given in the form X:D, where X is the chain length and D is the position of the double bond. The more negative the binding affinity, the more likely the chain is to bind.}
    \label{fig:affinity}
\end{figure}

\subsection{Molecular Dynamics}
\label{sec:MD}

Once the process of docking fatty acids of varying carbon length was completed, we proceeded to run molecular dynamics (MD) simulations on the protein as well as the docked protein-ligand complex. This process allows us to visualize which domains of the protein fluctuate over a given period of time. MD simulations afford us opportunities to better understand how not only the protein, but a specific protein-ligand pair evolves over time. More specifically, MD provides us information on protein allostery and protein docking \citep{Hertig2016}. This work is based around detailing the structure-function relationship of the FAD-class of proteins and is why a focus on docking and MD Simulations are the most critical aspects of this study. 

Following the docking procedures taken in Section \ref{sec:Docking}, two sets of docking experiments were set up and run for each model: 1. The protein without substrate and 2.The protein docked with the highest observed binding affinity ligand. Each of which were run for 100ns and motions were quantified with a root mean square fluctuation (RMSF) calculation. By utilizing this methodology, we were able to calculate which regions of the proteins move naturally, and which are most affected by the binding of a ligand within the binding pocket. As molecular dynamics solutions are computationally expensive, we selected one protein-ligand complex from the docking solutions for each protein model. We selected the palmitoleic acid (16:1)-AtFAD6 complex and oleic acid (18:1)-AtFAD2 complex. These FAs were chosen since they are the predominant binding candidates for their respective proteins. This selection gives us specifics into how the protein evolves over time when carrying out it's fundamental application in the cell. 

We plotted the resulting RMSF plots for all the cases run and overlaid them into a concise plot. Figure \ref{fig:rmsf} shows the color coded RMSF plots for AtFAD6 and AtFAD2 with and without the docked protein included. It is apparent that when the docked FA is included, the overall RMSF increases, but in specific regions, the increase is much more pronounced. For example, in AtFAD6 there is a major overall increase in the RMSF of the yellow region associated with a large portion of the cytosolic cap. This could indicate that the docking of a FA within that region alters the space available in the binding pocket. Another observation is that the links between trans-membrane helices contribute to much of the motion within the protein. The helices themselves are rather stationary over time which would make sense considering they are embedded into the lipid bi-layer. The same type of motion can be seen within the AtFAD2 protein as well. The inclusion of the docked FA increases the overall RMSF of the regions included in the cytosolic cap region of the protein and remain more static in the trans-membrane region. 

\begin{figure}[!ht]
    \centering
    \includegraphics[width=\textwidth]{rmsf_fixed.png}
    \caption{Root mean square fluctuation (RMSF) of backbone of AtFAD6 (top) and atFAD2 (bottom) without the substrate (solid line) and with the substrate (colored profile coding different structural regions). Palmitoleic Acid is the substrate for AtFAD6 and oleic acid is the substrate for AtFAD2. The red line in the AtFAD2 plot is at the break in residues not included in the homology model.}
    \label{fig:rmsf}
\end{figure} 

\subsection{Machine Learning Methods Used in Protein Classification and Function Prediction}

Machine learning introduces an effective way to classify proteins and further predict function. In this section we will examine some of the available options and how they can be useful for the analysis of Arabidopsis FADs. 

There are many acceptable current options for protein sequence/structure alignment and homology modeling. It is beneficial to be able to predict a protein's structure from its amino acid sequence therefore protein structure prediction has been a major focus in the community. The primary means is by aligning the target sequence with a number of homologous protein sequences, for some of which the structures (templates) are known, and thereby predicting the resulting three-dimensional structure of the target sequence. 

More recently, there have been large strides taken in the domain of machine learning for use in building relationships between protein sequences and their final characterized secondary, tertiary, and quaternary structures. The benefit to having a comprehensive machine learning model, is that simple single sequences can be quickly and more accurately characterized due to the detailed nature of machine learning models. These models are able to be trained on very comprehensive and well understood structure-function relationships and can precisely draw the necessary relationships required for alignment. 

Notable examples of machine learning applications towards protein structure prediction is
that of AlphaFold \citep{Jumper2021} and RoseTTAFold \citep{Baek2021}. The AlphaFold team
constructed a neural network trained on the three-dimensional
distances between amino acid residues with a ground-breaking
level of accuracy. The RoseTTAFold team implements a three-track network in which information at the one-dimensional (1D) sequence level, the 2D distance map level, and the 3D coordinate level is successively transformed and integrated. The application of these workflows pertaining to the current work was considered but ultimately not used as our existing structures shared a high enough structural similarity with AlphaFold/RoseTTAFold that we remained with our homology structure. Those comparisons can be found in the Supplemental Information.

\section{Conclusion}
Fatty acid desaturates are of great interest for basic understanding of how proteins act catalytically but also, for translational uses in  helping develop agricultural crops that are used for either biofuels or as chemical feedstocks.  Our current work is exclusively using computational approaches to predict the structure  and motion of the desaturases using software as  the pipeline and the great number of sequences available from many organisms.Predictions from such approaches can be empirically tested by designing and producing mutant proteins for analysis \citep{Cai2020,Cai2018}.

FADs present a critical research target within the area of sustainable platform chemical production due to the substrate and product diversity. Because lipid synthesis is highly demanding for both energy and reducing power, there is great interest in developing fundamental understanding of functionality regarding substrate specificity, regioselectivity, oxidation chemistry, and cofactors.

Activities of membrane-bound FADs in plants (and many organisms) are modulated by temperature perhaps by directly sensing their membrane lipid environments. Future experiments will extend these studies to investigate the effects of temperature and the membrane environment as factors that might influence FAD activity.  This is a minor criticism perhaps, but in the context of understanding all aspects of catalysis, these two parameters are important for many membrane-bound FADs.

\textbf{Future vision and Outlook-- Characterize}

Our present research is focused on the development of workflows that will be able to combine substrate docking and molecular dynamics for the characterization of fatty acid desaturase (FAD) enzymes. By being able to analyze how fatty acids of varying lengths bind within the diiron enzyme active sites of FADs, we can find ways to elucidate binding specificity and overall activity. Informed computational analysis reduces the time spent experimentally identifying and testing combinations of FADs with fatty acids and allows for more application-focused work.

\section{Methods}
\label{sec:results}

\subsection{FAD Curation}
The phylogenic tree diagram and multiple sequence alignment of the 10 Arabidopsis FADs displayed in Section 
\ref{sec:Curate} were generated using Clustal$\Omega$ \citep{Madeira2019}. It is a web-based server with the 
only input being a file containing each FASTA format sequence from each protein.

\subsection{Homology Modeling}
The PHYRE2 \citep{Kelley2015} application was used to generate an optimized homology model from a variety of 
known template structures. We selected pdb:4ZYO as the template model and the primary sequence FASTA files 
of the two target proteins (AtFAD6, AtFAD2). This application was utilized with it's web-based server.

\subsection{Docking}
Docking simulations were run through QVINA \citep{Hassan2017} with a selection of substrates. The docking 
region was refined from a blind dock to a targeted dock in the region surrounding the diiron core and inside 
the binding pocket using AutoDockTools.

The optimization of the pre-docked FA chains was done using NWChem 6.8.1.30 \citep{Apra2020}. 
Partial charges of the FA chains were obtained based on the restrained electrostatic potential atomic partial charge (RESP) 
method \citep{Bayly1993} at the HF/6-31G* level. 

\subsection{Molecular Dynamics}
\label{sec:Molecular}
Molecular dynamics simulations were carried out using a 
GROMACS \citep{Lindahl2021,Hess2008,Miyamoto1992} workflow with the homology models AtFAD2 and AtFAD6.  
The top docking candidates for each protein were taken and used as the initial poses for the protein-ligand complex 
simulations. The Amber FF14SB force field \citep{Maier2015} was used for the 
protein along with the general AMBER force field (GAFF) \citep{Wang2004}.
The TIP3P water model \citep{Jorgensen1983} with Joung and Cheatham ion parameters \citep{Joung2008} were used for 
establishing a cubic box of water and neutralizing monovalent ions in the system. 
The initial system was minimized by the conjugated gradient algorithms up to a maximum residual force of 10.0 KJ/mol. The 
system was then equilibrated at 300K for 500 ps under NVT ensemble using the 
modified Berendsen thermostat velocity rescaling method \citep{Bussi2007},
followed by 500ps with NPT ensemble using the Berendsen pressure coupling method \citep{Berendsen1984}. 
Harmonic restraints with force constant of 1,000 KJ/mol were applied on the 
protein and substrate during equilibration steps. After the equilibrations, we performed 
production runs for 100 ns at 300 K and 1 atm using the 
Parrinello-Rahman pressure coupling method \citep{Parrinello1981,Nose1983}. All  of the 
restraints were released during this step. The timestep adopted in all of the simulations are 2 fs and a particle mesh Ewald 
algorithm \citep{Essmann1995} was used to evaluate long-range electrostatic interactions. RMSF calculations on the results were also carried out using the GROMACS rmsf calculation tool.


\subsection{Visualizations}
All the protein-ligand visualizations done within this work were made using PyMOL \citep{PyMOL}.

\section*{Declaration of Competing Interest}
The authors declare that they have no known competing financial interests or personal relationships that could have appeared to influence the work reported in this paper.

\section*{Acknowledgements}
This work was supported in part by the U.S. Department of Energy, Office of Science, Laboratory Directed Research Funding (LDRD), at the Pacific Northwest National Laboratory. Pacific Northwest National Laboratory (PNNL) is a multiprogram national laboratory operated by Battelle for the DOE under Contract DE-AC05-76RLO 1830. J.S. was supported by the Physical Biosciences Program, and Photochemistry and Biochemistry group within the US Department of Energy (DOE), Office of Science, Office of Basic Energy Sciences, Division of Chemical Sciences, Geosciences and Biosciences (grant KC0304000). This research used computational resources provided by Research Computing at the Pacific Northwest National Laboratory.
%% The Appendices part is started with the command \appendix;
%% appendix sections are then done as normal sections
\appendix

\section{AlphaFold Similarities}
\label{sec:sample:appendix}

To compare our two homology models with the newly released AlphaFold structures, we compare the structural and sequence similarity between the proteins. We primarily look at the high confidence regions that are predicted by AlphaFold. The tools used for comparing structures was the PDB Structural Alignment Tool with jFATCAT \citep{Ye2003,Li2020} \& JCE \citep{Shindyalov1998} and the DALI protein structure comparison server \citep{Holm2020}. Overall the AtFAD6 protein contained more similarities to the AlphaFold structure than AtFAD2. The specific numbers can be found in the \ref{table:metric}. We concluded that even though the homology models don't perfectly align with the AlphaFold structures, they are similar enough to utilize in this work.

\begin{table}[!ht]
\begin{center}
\caption{A full collection of comparison metrics explored in comparing the homology models with the AlphaFold and RoseTTAFold structures. In each case the protein listed is compared to the homology model. AF-FAD is the AlphaFold modeled protein compared to the homology model and RF-FAD is the RoseTTAFold modeled protein compared to the homology model.}
\begin{tabular}{lrrrrr}
\hline
 & DALI &  &  &  & \\
\hline
Protein & rmsd & Z-Score & Length & \%id & \\
AF-FAD2 & 4.2 & 8.8 & 181 & 43 & \\
AF-FAD6 & 4.3 & 7.8 & 183 & 49 & \\
RF-FAD2 & 5.0 & 7.4 & 189 & 38 & \\
RF-FAD6 & 4.2 & 8.7 & 191 & 50 & \\
\hline
 & FATCAT &  &  &  & \\
\hline
 & rmsd & Score & Length & SI\% & SS\%\\
AF-FAD2 & 4.17 & 328.95 & 227 & 33 & 38\\
AF-FAD6 & 3.29 & 327.83 & 217 & 60 & 66\\
RF-FAD2 & 4.3 & 317.04 & 215 & 48 & 53\\
RF-FAD6 & 3.5 & 325.17 & 229 & 61 & 66\\
\hline
 & JCE &  &  &  & \\
\hline
 & rmsd & Score & Length & SI\% & SS\%\\
AF-FAD2 & 4.26 & 502.85 & 190 & 37 & 43\\
AF-FAD6 & 3.98 & 597.53 & 176 & 49 & 57\\
RF-FAD2 & 4.52 & 449.33 & 191 & 29 & 37\\
RF-FAD6 & 4.37 & 448.89 & 206 & 48 & 54\\
\end{tabular}
\end{center}
\label{table:metric}
\end{table}

\subsection{Sequence Similarity}

The first comparisons we can make are between the sequence similarities. By visualizing the overlapping region between the homology model and the AlphaFold structure, we look at which sequences are conserved between the two. Represented in \ref{fig:Con_Seq} obtained from the DALI server. The overall sequence similarity is relatively consistent between our models, and each structure has similarities in key regions of the protein transmembrane region and cytosolic cap. The AtFAD6 protein has a higher sequence similarity than AtFAD2 overall between the different methods outlined in \ref{fig:Con_Seq}.

\begin{figure}[!ht]
    \centering
    \includegraphics[width=0.5\textwidth]{Omega6_Conserved_AF.png}\hfill
    \includegraphics[width=0.5\textwidth]{Delta12_Conserved_AF.png}\hfill
    \caption{Sequence similarity between homology models and AlphaFold structures. The left figure represents the conserved sequence region within the AtFAD6 and AlphaFold protein and the right figure represents the AtFAD2 and it's AlphaFold structure. The blue regions are the regions with perfect alignment while the green and yellow represent lessining similarity.}
    \label{fig:Con_Seq}
\end{figure}

\subsection{Structure Similarity}

The next comparison to make between the homology models and AlphaFold is the structural similarity. This is an important analysis to make as it can have a significant impact on the molecular dynamics simulation. The sequence similarity visual representation provided by DALI is shown in \ref{fig:Con_Struc}.

\begin{figure}[!ht]
    \centering
    \includegraphics[width=0.5\textwidth]{Omega6_Con_Structure_AF.png}\hfill
    \includegraphics[width=0.5\textwidth]{Delta12_Con_Structure_AF.png}\hfill
    \caption{Caption}
    \label{fig:Con_Struc}
\end{figure}

\section{FAD Reaction Mechanism}

To better understand how different FADs chemically add a
double bond to a fatty acid chain, we need to understand the
processes involved as well as the different types of
mechanisms. Each desaturase pathway starts with reduced
nicotinamide adenine dinucleotide (phosphate) (NAD(P)H),
followed by an electron transfer by a flavin adenine
dinucleotide containing reductase either to cytochrome b5
(cyt\textit{b}5) or ferredoxin (Fd), which bind with the desaturase for
further use in the activation of the diiron core \citep{Shanklin1998}. From
here, the process is similar in that desaturation of the fatty
acid occurs, but the location in which it occurs is separate
and where the desaturase obtains its electrons is specific to
$\Delta$ / $\omega$-desaturases. 

\begin{figure}[!ht]
    \includegraphics[width=\textwidth]{Reac_Path.png}
    \caption{Overview of the linoleate biosynthesis I pathway in Arabidopsis thaliana 
    (AraCyc). This pathway illustrates the desaturation of a C18:1 fatty acid (oleic 
    acid) into a C18:2 fatty acid (linoleic acid). Plant FAD tend to act on 
    glycerolipids over acyl-CoA structures, so the beginning and end of the pathway 
    contain some lipid transferase to convert an acyl-CoA with a 
    fatty acid sidechain to a glycerolipid with a fatty acid 
    sidechain and vice versa.}
    \label{fig:linoleate}
\end{figure}

\subsection{$\Delta$-desaturases}

The $\Delta$-desaturases work by appending a
double bond to a fatty acid from the front-end or carboxyl
end. Figure \ref{fig:fad2} shows the FAD2 reaction in the linoleate
biosynthesis I pathway (Complete pathway shown as Fig \ref{fig:linoleate}). It
is a major enzyme responsible for the synthesis of 18:2 fatty
acids in the endoplasmic reticulum. It contains His-rich
motifs, which contribute to the interaction with the electron
donor cyt\textit{b}5 \citep{Berardini2015}. FAD2 introduces a second double bond in
linoleate in linoleate-containing phosphatidylcholine, an
important extra-plastidial membrane lipid \citep{Covello1996,Petrini2004,Caspi2012}.

\begin{figure}[!ht]
    \includegraphics[width=\textwidth]{Picture2.png}
    \caption{Overview of the FAD2 $\Delta$12 desaturase reaction (E.C 1.14.19.22). This 
    reaction takes a fatty acid side chain oleic acid attached to a glycerophospholipid 
    moiety and acts to increase its chain length by one to a linoleic acid sidechain 
    with a glycerophospholipid moiety. This reaction requires the donation of electrons 
    from a cytochrome b5 molecule along with molecular oxygen.}
    \label{fig:fad2}
\end{figure}

It was suggested and shown that cytochrome b5 is involved in
the desaturation of fatty acids in the endoplasmic reticulum
as an electron donor \citep{KEARNS1991431,Smith1990}. This process subverts the need
for the activation of the diiron core, as the electrons are
donated from cyt\textit{b}5. 

\subsection{$\omega$-desaturases}

\begin{figure}[!ht]
    \includegraphics[width=\textwidth]{fad6.png}
    \caption{Overview of the FAD6 catalyzed omega 6 fatty acid desaturation reaction 
    (E.C. 1.14.19.23). Similar to the $\Delta$12 desaturation reaction, this pathway 
    transfers a C18:1 oleic acid to a C18:2 linoleic acid. The main difference is the 
    source of the donated electrons coming from a ferredoxin diiron core at the active 
    site with molecular oxygen for activation instead of binding with cyt\textit{b}5.}
    \label{fig:fad6}
\end{figure}

In contrast to the microsomal $\Delta$-desaturases, FADs in
chloroplasts involve Ferredoxin (Fd), Fd-NADP+ oxidoreductase,
and NADPH, and which suggests that Fd is the potential
candidate for being the donor of electrons to chloroplast
desaturases \citep{Schmidt9477,Schmidt1990}. This type of desaturase is also referred
to as an $\omega$-desaturase, as it works to append the double
bond on the methyl end of the fatty acid chain. This
$\omega$-desaturase pathway is shown on Fig X as E.C.
1.14.19.23 which represents the Omega-6 fatty acid desaturase
found in the chloroplast. Fig \ref{fig:fad6} shows a more detailed view of
the FAD6 reaction. This plastidial enzyme is able to insert a
cis double bond in monounsaturated fatty acids incorporated
into glycerolipids. The enzyme introduces the new bond at a
position 3 carbons away from the existing double bond, towards
the methyl end of the fatty acid \citep{Falcone1994,Mekhedov2000}. It is primarily
responsible for the synthesis of 16:2 and 18:2 fatty acids
from galactolipids, sulpholipids and phosphatidylglycerol\citep{Berardini2015}.


%% If you have bibdatabase file and want bibtex to generate the
%% bibitems, please use
%%
\bibliographystyle{elsarticle-num-names} 
\bibliography{paper}

%% else use the following coding to input the bibitems directly in the
%% TeX file.

%\begin{thebibliography}{00}

% %% \bibitem[Author(year)]{label}
% %% Text of bibliographic item

% \bibitem[ ()]{}

%\end{thebibliography}

\end{document}

\endinput
%%
%% End of file `elsarticle-template-num-names.tex'.
