## Supplementary figures and images for "Data-Driven Modeling and Analysis of Fatty Acid Desaturase in Plants"

### Binding_Channels.png

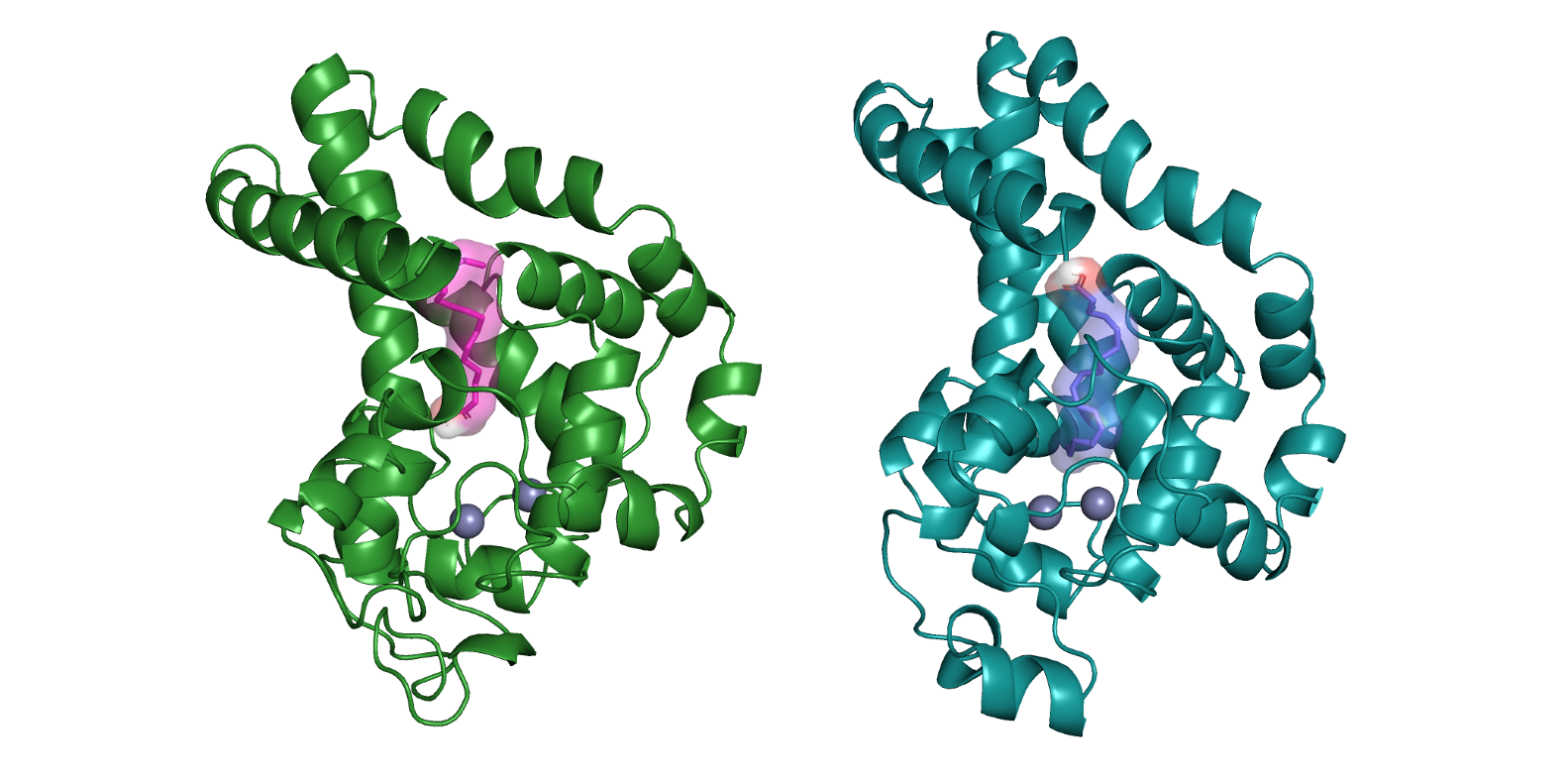

### Delta12_Con_Structure_AF.png

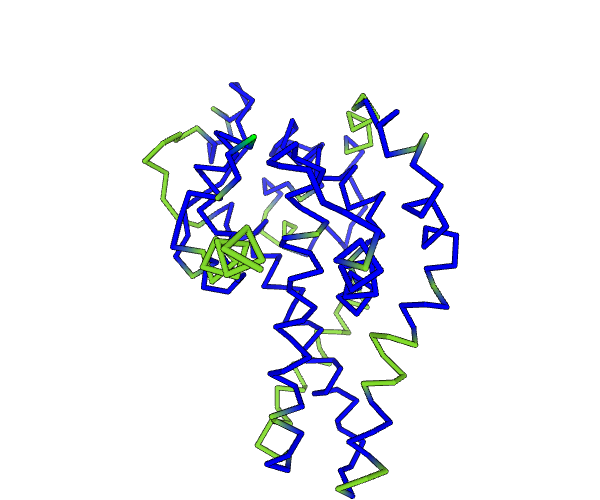

### Delta12_Conserved_AF.png

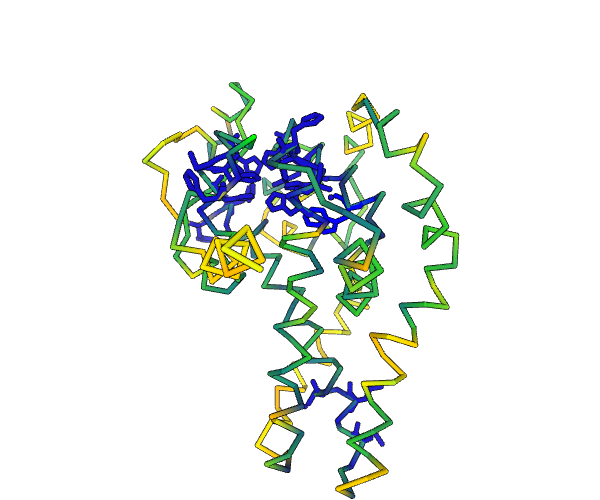

### fad6.png

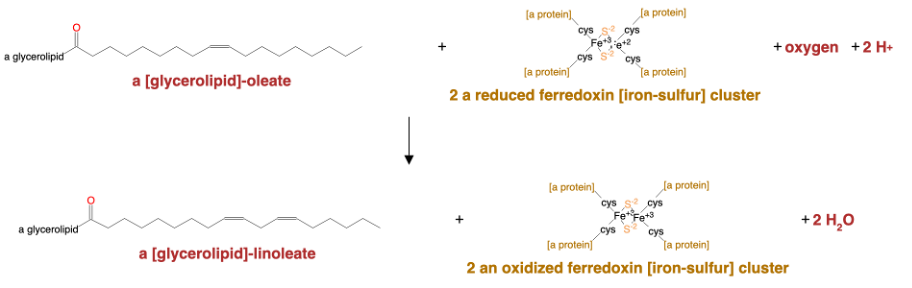

### FAD_BA_STACK.png

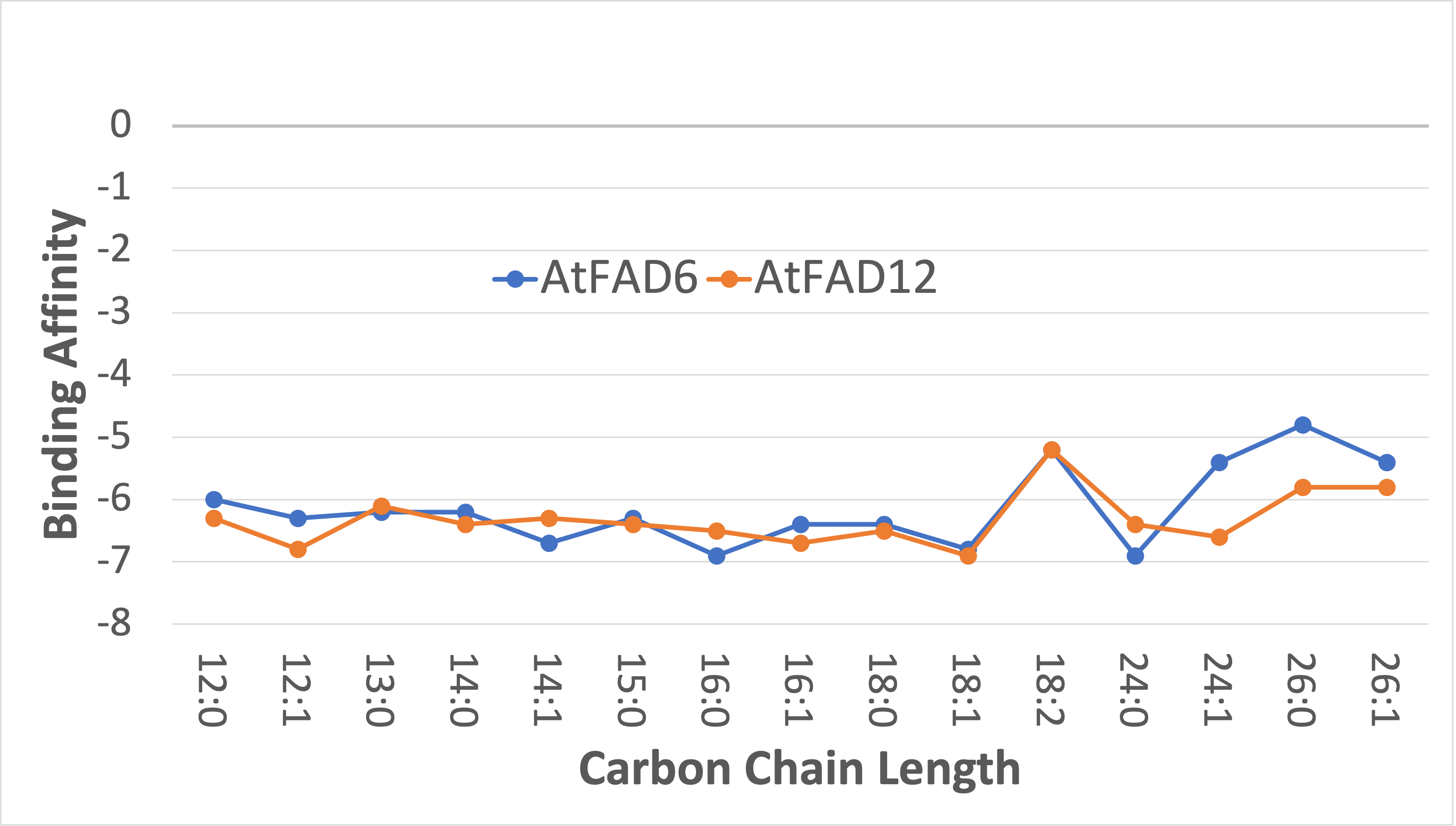

### FAD_GRAPHICAL_ABSTRACT.png

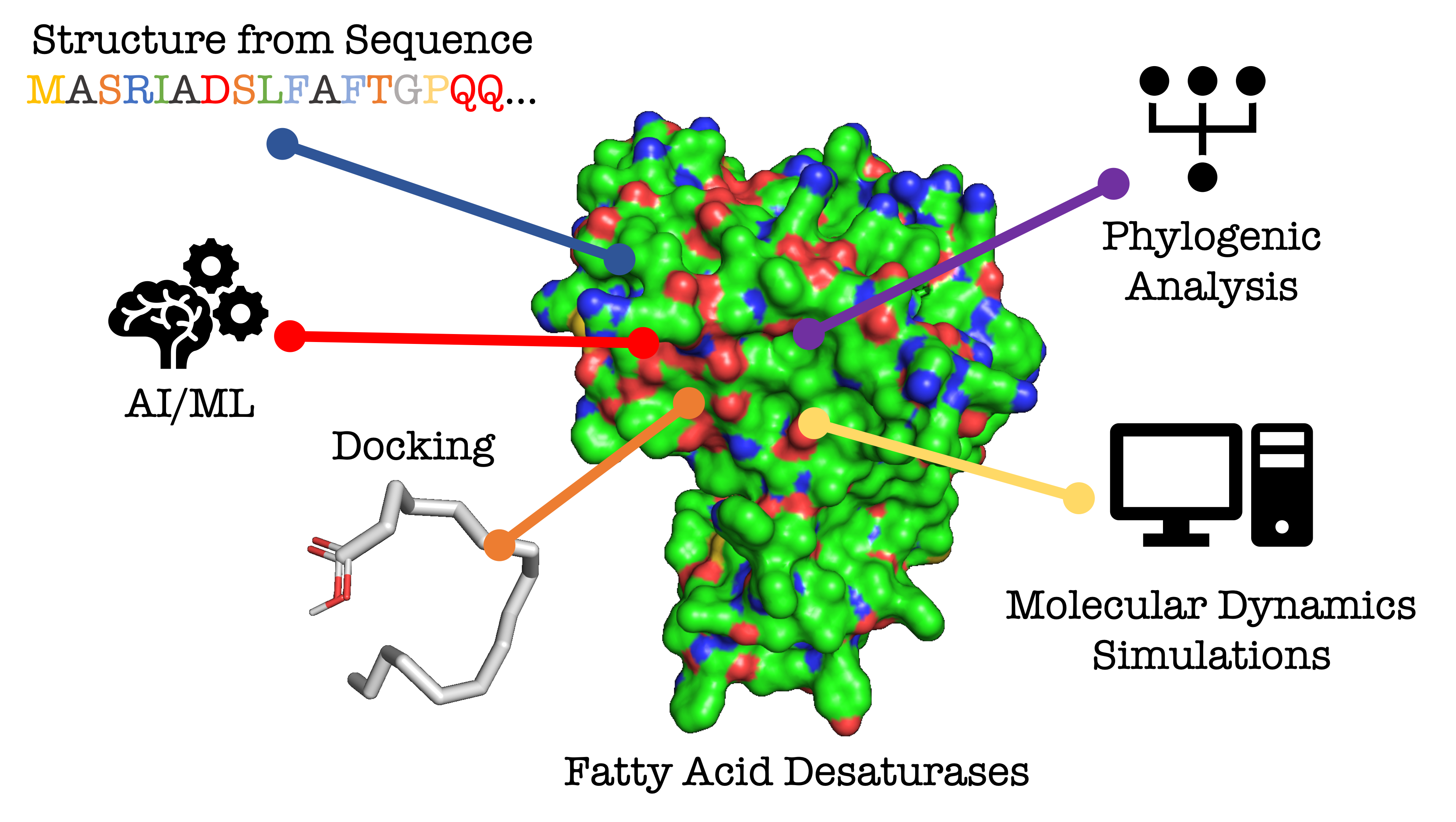

### Hist_Regions.png

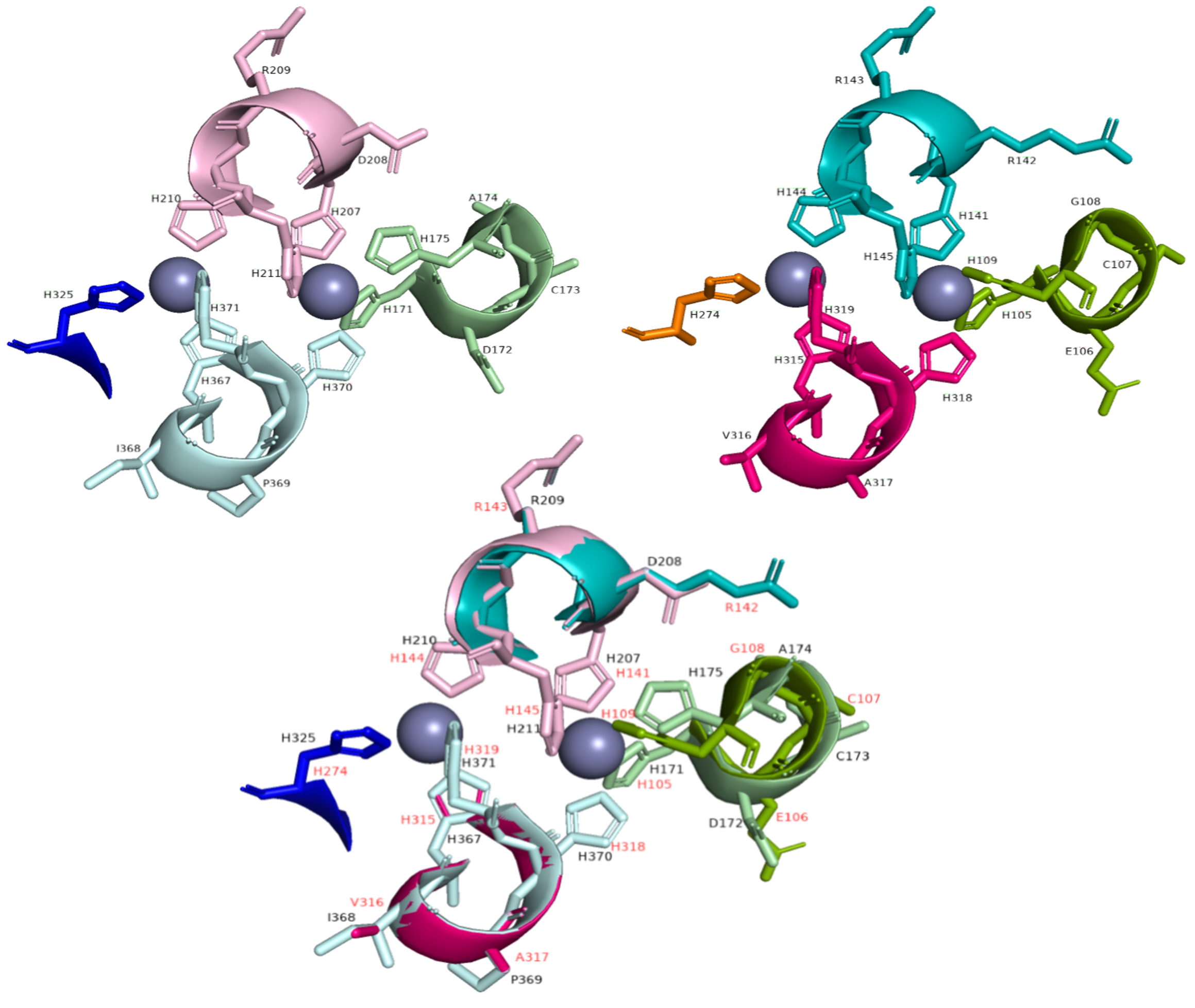

### Homology_Full.png

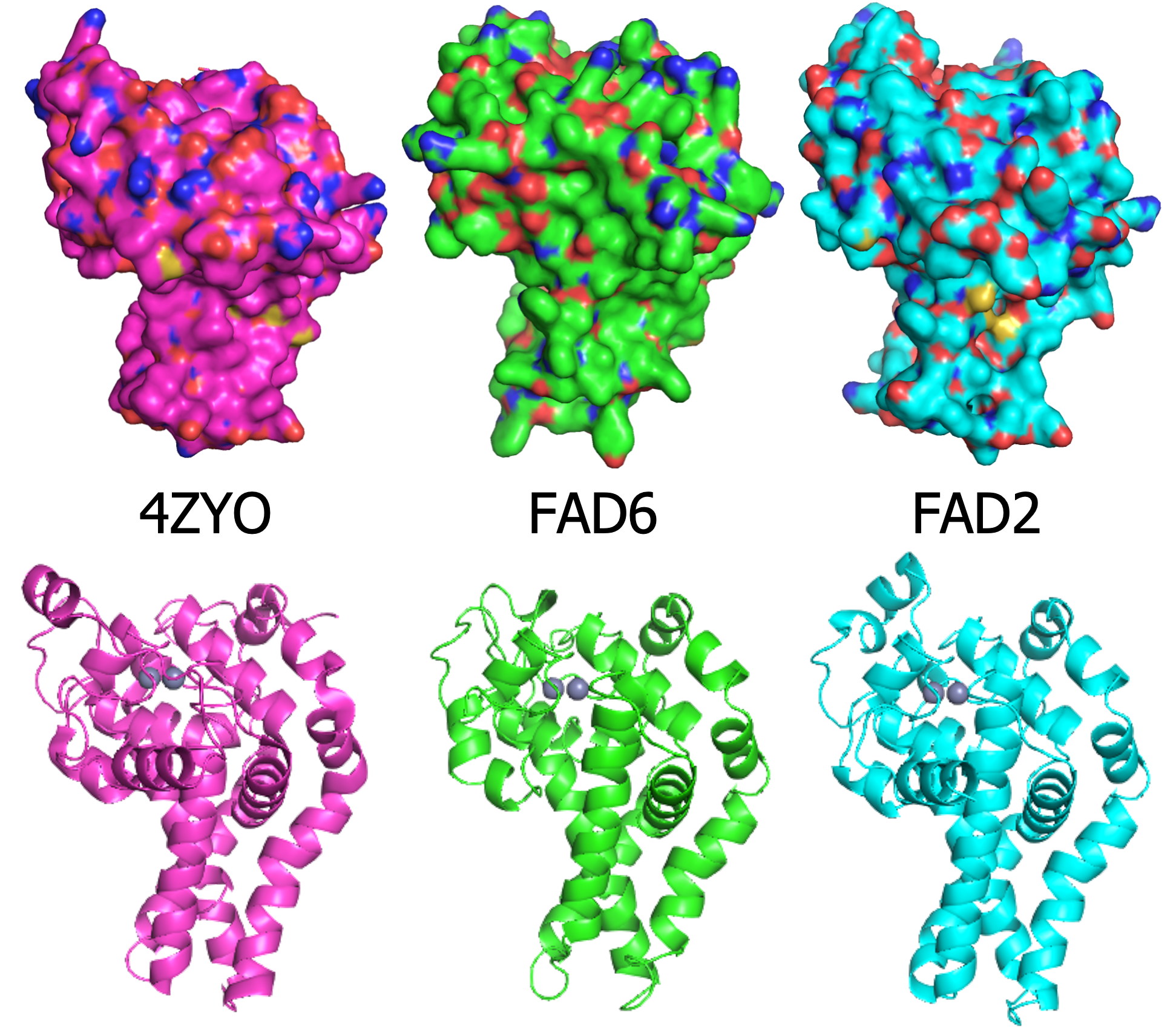

### Omega6_Con_Structure_AF.png

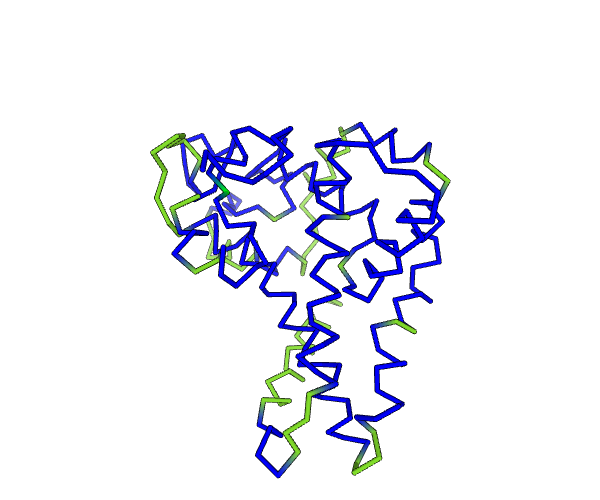

### Omega6_Conserved_AF.png

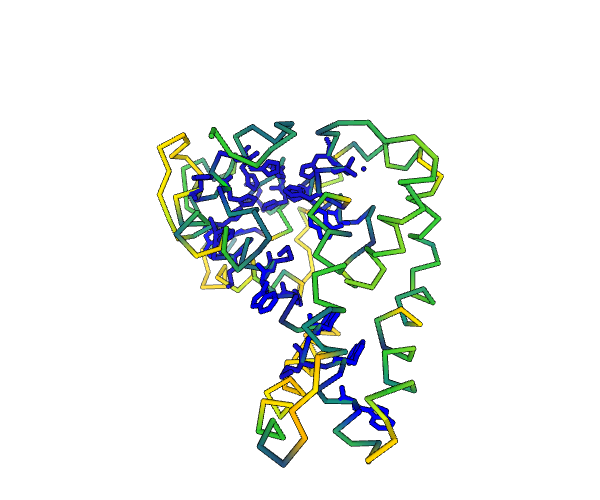

### phylo.png

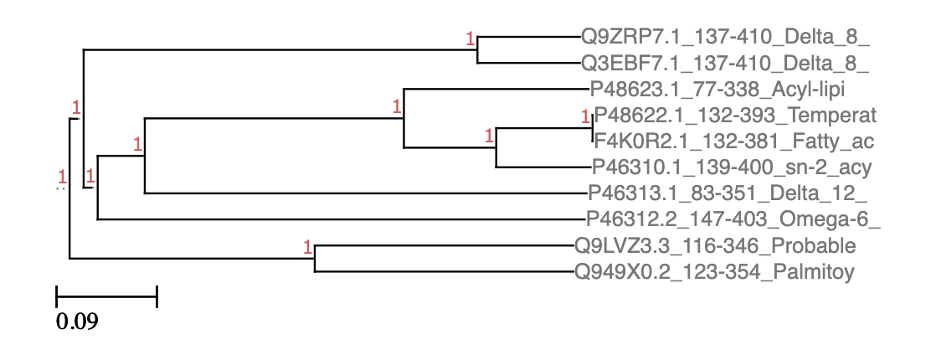

### Picture2.png

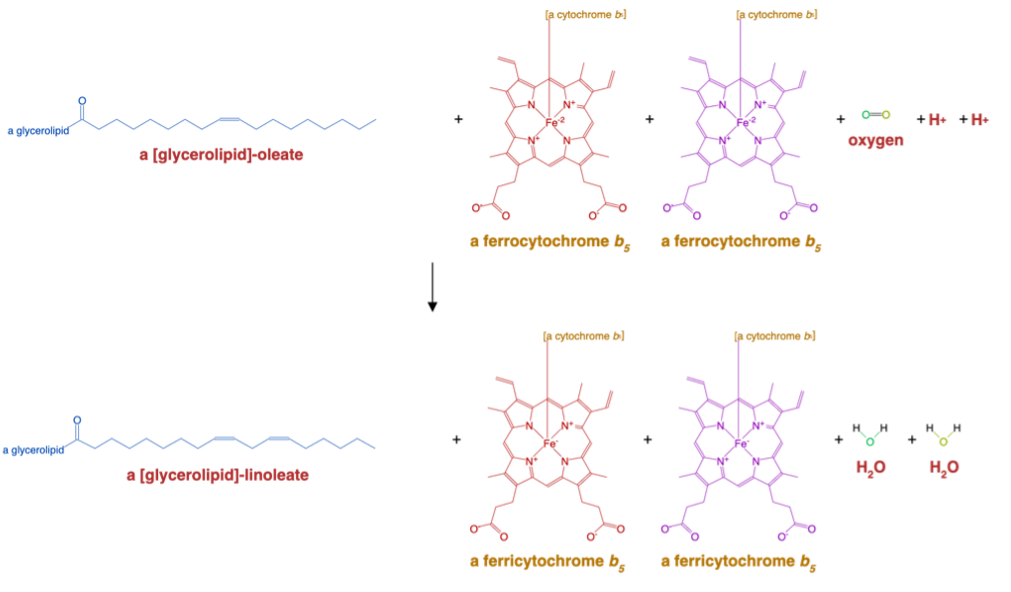

### Reac_Path.png

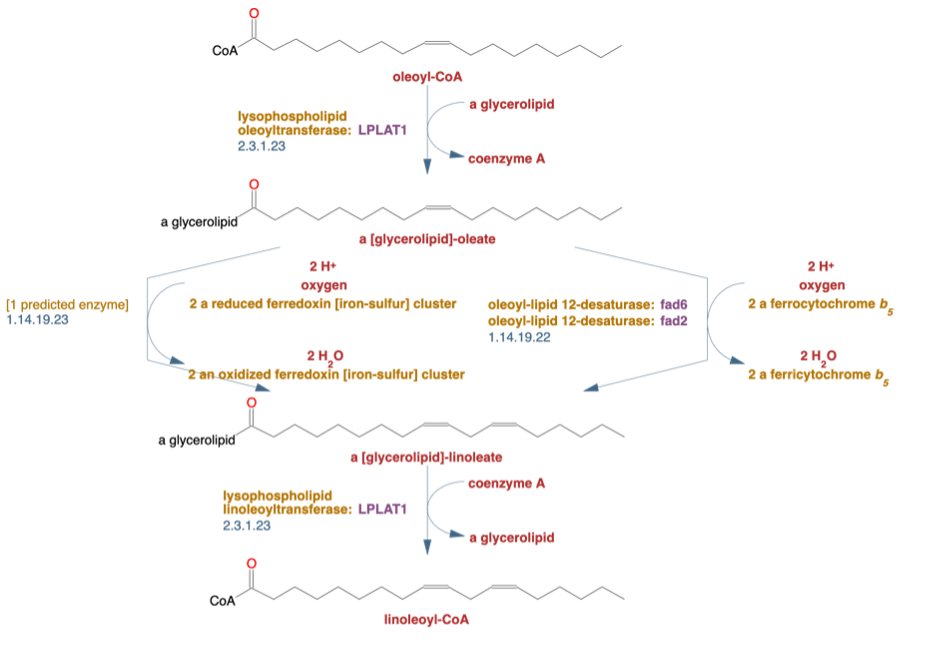

### rmsf_fixed.png

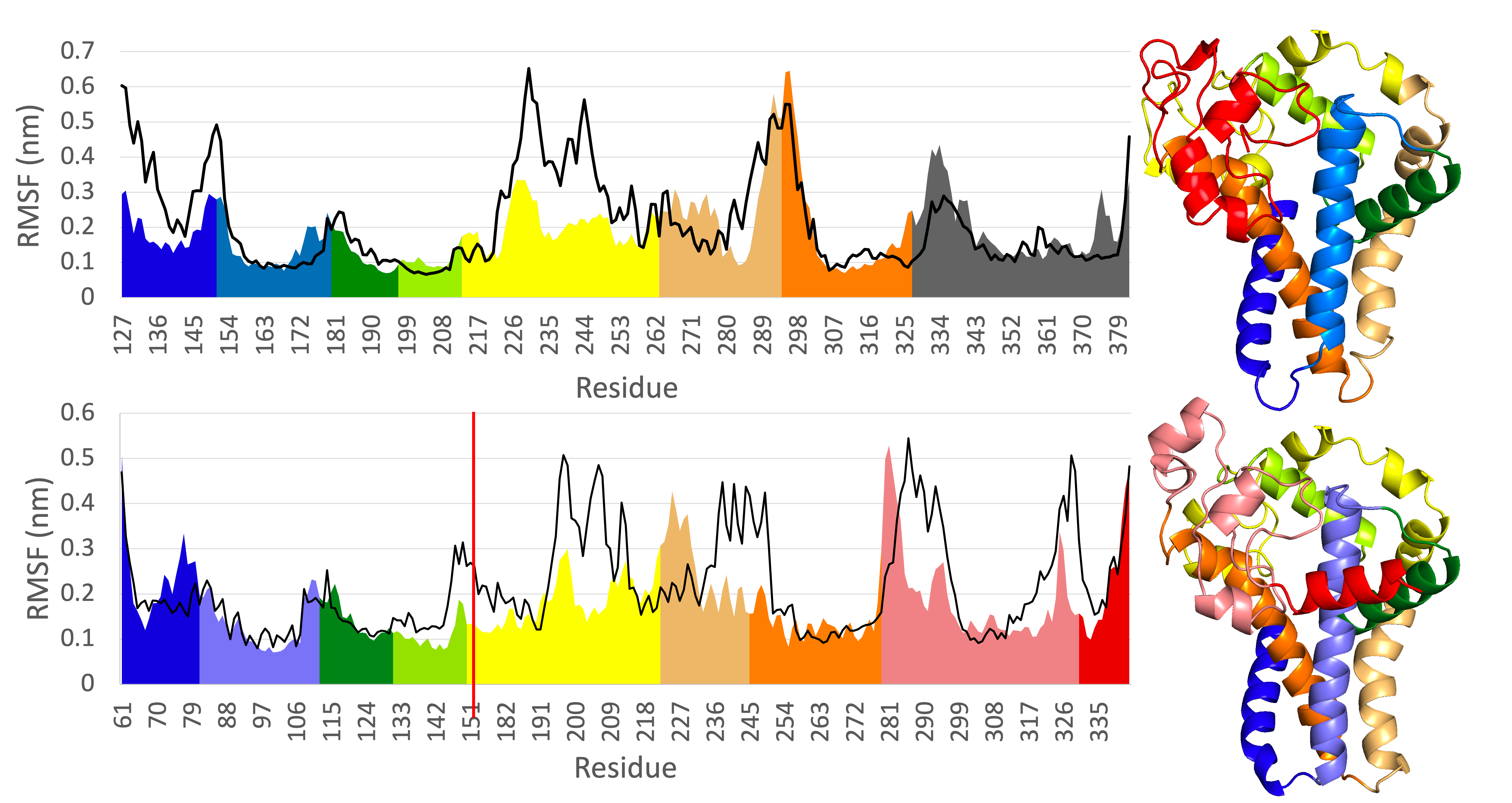

### Scheme.png

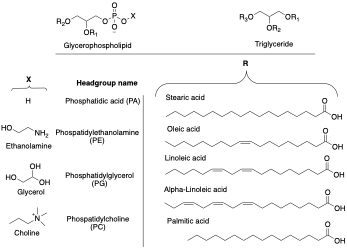

### Screen Shot 2021-03-18 at 3.23.26 PM.png

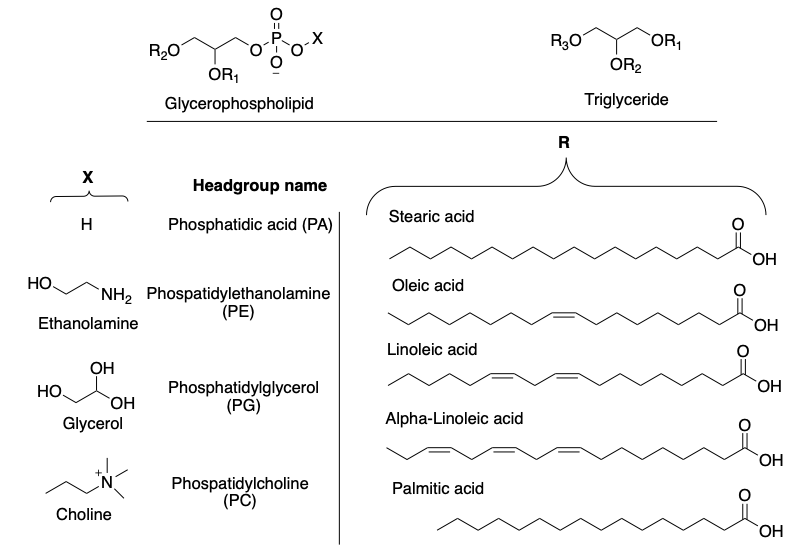

### Sequence_Box.png

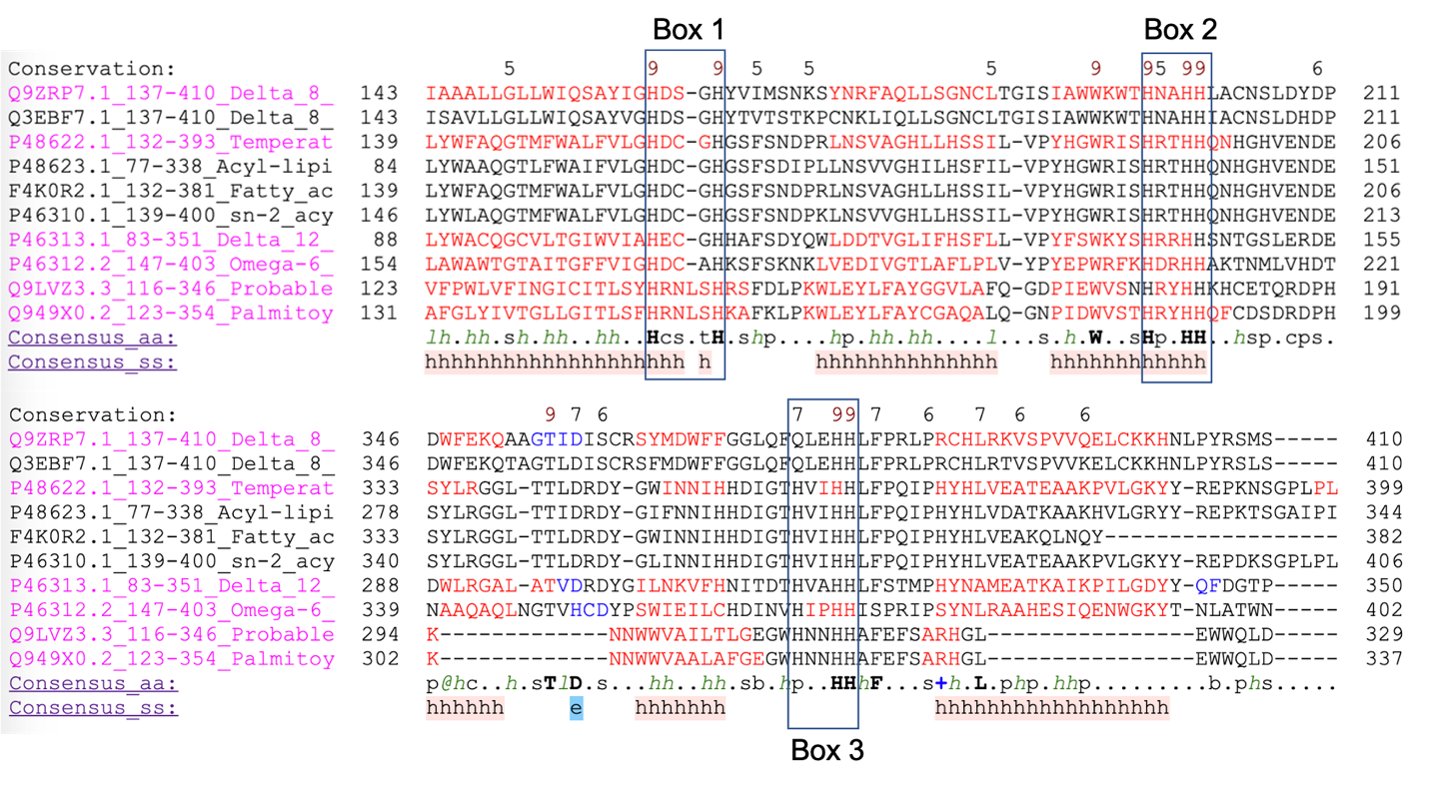

### three_proteins.png

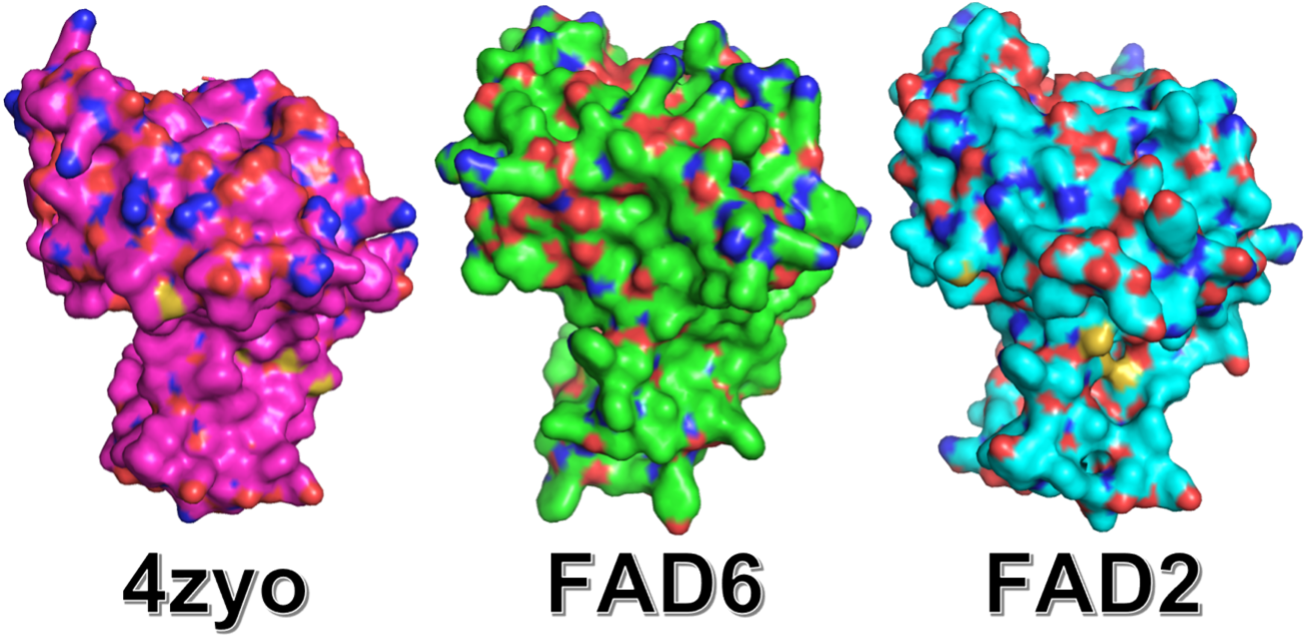
